# NUP62 acts as T cell activation-dependent switch to regulate post nuclear import steps of HIV-1 replication in primary CD4 T cells

**DOI:** 10.64898/2026.11.26.768590

**Authors:** Swetha Ananth, Ina Ambiel, Samy Sid Ahmed, Carlotta Penzo, Bojana Lucic, Kristian Bodulić, Laetitia La Rocca, Yeonsu Kim, Fariha Mahzabin Annesha, Barbara Müller, Venera Weinhardt, Oliver T. Keppler, Hans-Georg Kräusslich, Marina Lusic, Oliver T. Fackler

## Abstract

Resting CD4 T cells resist productive HIV-1 infection. T cell activation licenses their permissivity for infection, increases nuclear pore complex (NPC) density, and alters chromatin organization. Paradoxically, HIV-1 nuclear import is more efficient in resting than in activated CD4 T cells, despite lower NPC abundance. Using activation-neutral gene editing in primary human CD4 T cells, we show that T cell activation reshapes nucleoporin (NUP) contributions to infection. The FG-nucleoporin NUP62 switches from an inhibitor of HIV infection in resting cells to a dependency factor in activated cells. In resting cells, NUP62 depletion modestly increases nuclear import, shifts proviral integration away from transcriptionally active nuclear regions, and enhances viral transcript abundance and gene expression. In activated cells, NUP62 depletion leaves steps through integration unaffected but strongly reduces nascent viral transcripts and gene expression. T cell activation switches previously unrecognized post-nuclear-import functions of NUP62, uncoupling early and late steps of HIV-1 replication.

## Introduction

Viral infections typically follow an organizing principle in which there is a stepwise reduction in the amount of infectious substance, and the efficacy of each step serves as an important parameter for subsequent steps. Barriers that reduce infectious particles include natural physical barriers, the availability of cellular resources required for virus replication, and the antiviral activity of host cell innate immune factors (Harris et al. 2012, Pan et al. 2013, Simon et al. 2015). The magnitude by which individual steps reduce infection rates is an important characteristic of individual virus - target cell systems.

For HIV-1, the nuclear envelope constitutes an important natural host cell barrier that needs to be overcome to import viral genomes into the nucleus for subsequent integration into the host cell genome. To achieve this, the capsid protein (CA), assembled into cone-shaped capsids containing the subviral genomic complex, acts as a karyopherin and is the central viral determinant of nuclear import by active transport through nuclear pore complexes (NPCs). NPCs represent large protein assemblies of approximately 35 different nucleoporins (NUPs) that govern selective nucleocytoplasmic transport (Petrovic et al. 2026). Scaffold phenylalanine-glycine (FG)-NUPs such as NUP358 and NUP153 contribute to the architecture of the NPC, whereas others such as NUP98, NUP62 and NUP35 form a disordered mesh in the center through which bulky cargoes are selectively transported. Not all NUPs are stationary at the NPC, and they can exhibit varying residence times depending on cellular activation and proliferation states, resulting in high NPC heterogeneity that alters adaptor and regulatory functions (Rabut et al. 2004). Following a paradigm shift in the field driven by direct visualization of viral capsids by high-resolution microscopy, it is now well established that intact or nearly intact capsids are transported through the NPC (Zila et al. 2021, Shen et al. 2023, Dickson et al. 2024, Fu et al. 2024, Kreysing et al. 2025), which requires NPC elasticity and absorption of physical stress by the capsid (Hou et al. 2025, Kreysing et al. 2025). Furthermore, reverse transcription of the RNA genome into DNA is completed in the nucleus, where genome uncoating occurs shortly before integration (Burdick et al. 2020, Dharan et al. 2020, Selyutina et al. 2020). Capsids engage host cofactors such as cyclophilin A, CPSF6, and select NUPs to coordinate nuclear entry (Zila et al. 2021). Consistent with the central role of the NPC for capsid import, siRNA perturbations of NPC components, including NUP358, NUP153, NUP98, or NUP62 markedly reduce HIV-1 infection in model cell lines (Brass et al. 2008, Konig et al. 2008, Lee et al. 2010, Di Nunzio et al. 2012, Di Nunzio et al. 2013, Kane et al. 2018, Xue et al. 2023).

CD4 T lymphocytes, the most abundant HIV target cells in vivo, are particularly interesting to study host cell barriers since T cell activation, a physiological differentiation signal, completely alters their permissivity to infection. Resting cells that have not recently been stimulated by specific antigen are largely refractory to productive infection due to blocks to virus entry (limited dynamics of cortical and cytoplasmic actin filaments (Yoder et al. 2008, He et al. 2025), reverse transcription (reduced intracellular deoxynucleoside triphosphates (dNTP) pool due to dNTP degradation by the triphosphohydrolase SAMHD1) (Baldauf et al. 2012, Descours et al. 2012), and viral gene expression (low availability of cellular transcription factors (Budhiraja et al. 2013). T cell activation overcomes these barriers and allows CD4 T cells to promote all steps of the viral life cycle (Pan et al. 2013). Importantly, T cell activation also triggers fundamental changes in cellular metabolism and organization, including nuclear envelope remodeling and changes in chromatin architecture, which may impact HIV capsid nuclear import and/or integration (Smith-Garvin et al. 2009, Rawlings et al. 2011, Iqbal et al. 2021). Low levels of circularized viral DNA (vDNA) products existing as 2-LTR circles were previously interpreted as an indicator of low nuclear import efficacy in resting CD4 T cells (Baldauf et al. 2017). However, consistent with the finding that reverse transcription is completed in the nucleus, capsid visualization allowed us recently to establish that viral nuclear import is efficient in resting CD4 T cells (Ananth et al. 2024). This finding was unexpected as resting CD4 T cells contain relatively few NPCs, whereas T cell activation increases NPC abundance (Maul et al. 1971) and regulates the nuclear transport of cellular proteins (Shapiro et al. 2005). This raises the question of how HIV-1 adjusts its post-entry replication strategy to the changes in nuclear envelope and chromatin architecture of resting CD4 T cells.

Addressing this question requires the manipulation of individual cellular components in resting cells without compromising their resting state. However, until recently, gene editing of CD4 T cells was only possible following cell activation (Prelli Bozzo et al. 2024). Our recently developed highly efficient activation-neutral gene editing approach for resting CD4 T cells overcomes this technical barrier (Albanese et al. 2022, Morath et al. 2024). Here, we applied this approach to systematically examine how T cell activation modulates the NUP landscape in primary human CD4 T cells and to define the requirement for individual NUPs in HIV-1 post-entry replication. The results unexpectedly reveal NUP62 as a T cell activation-responsive molecular switch that governs HIV-1 replication downstream of nuclear import in CD4 T cells.

## Results

### T cell activation modulates HIV-1 nuclear import and infection in CD4 T cells

T cell activation induces extensive cellular remodeling to support robust effector functions (Smith-Garvin et al. 2009). We visualized primary human CD4 T cells isolated from buffy coats that were either kept resting or activated with anti-CD3/28 beads for 72 h by soft X-ray tomography and confocal microscopy (Fig 1A). T cell activation increased cells’ cytoplasmic-to-nuclear volume ratio and induced differences in X-ray absorption within the nucleus that can be attributed to changes in chromatin organization (indicated in 3D rendering of hetero-(blue) and eu-(green) chromatin in Fig 1A). Linear absorption coefficient curves in the nucleus showed that the average X-ray attenuation was reduced in activated CD4 T cells compared to resting conditions, highlighting chromatin decompaction (Le Gros et al. 2016) (Suppl. Fig. 1A). In addition, immunofluorescence analysis revealed an increase in the number of NUP98 puncta in the nuclear envelope of activated CD4 T cells, indicating elevated NPC density relative to resting CD4 T cells (Fig. 1A) (Maul et al. 1971).

**Figure 1.**
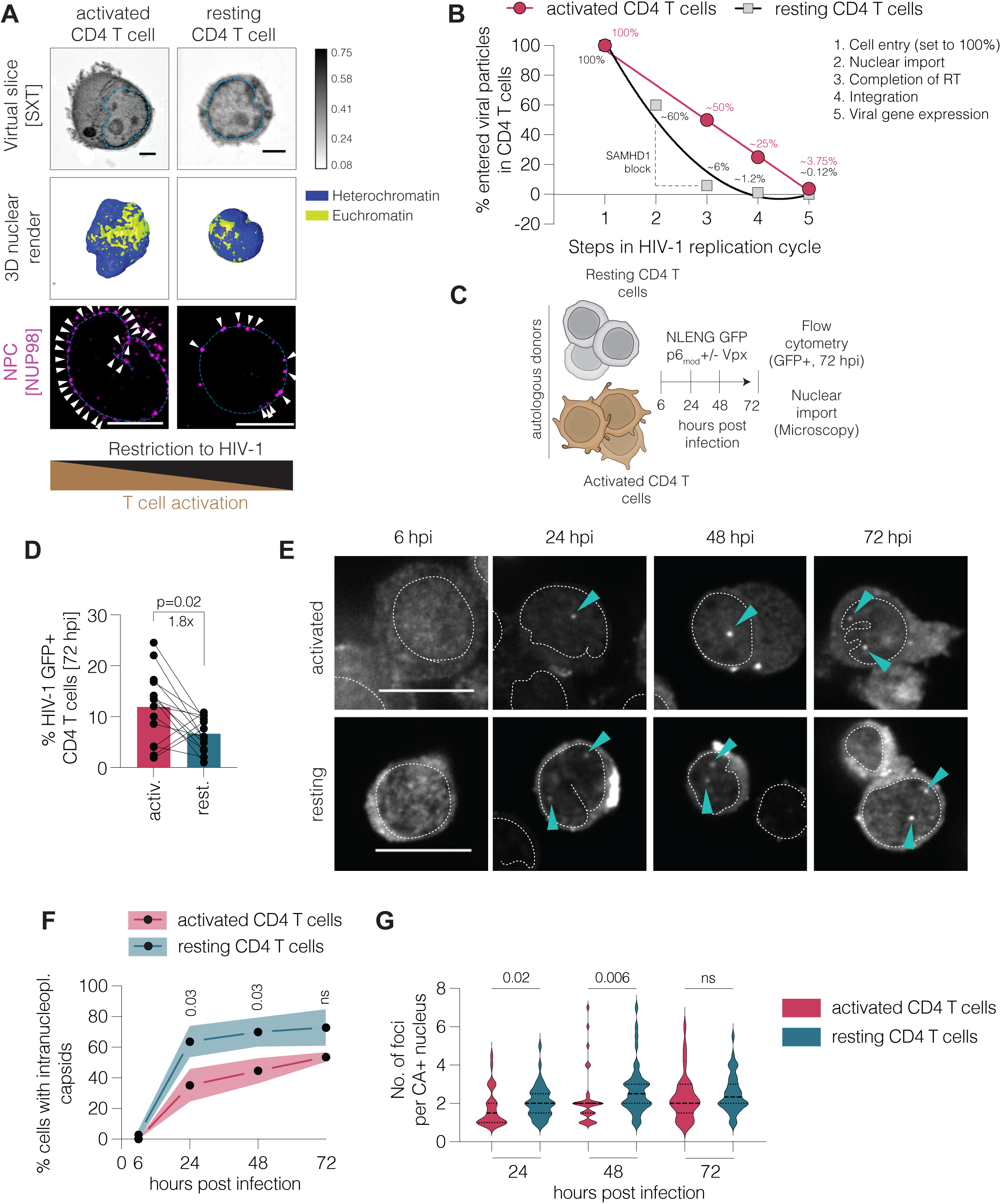
T cell activation alters HIV-1 infection in CD4 T cells. **(A)** Top: Single virtual slice reconstruction from soft X-ray tomography data obtained from peripheral CD4 T cells (resting and anti-CD3/CD28-activated). Dotted blue line highlights the nuclear boundary. Scale bar: 2 μm. Middle: 3D rendering of the nucleus from the cells shown above highlighting heterochromatin and euchromatin regions (in blue and yellow respectively) based on differences in linear absorption coefficients. Below: Representative Airyscan images of NPC distribution, visualized by NUP98 staining, in activated and resting CD4 T cells. Arrows indicate NUP98 puncta overlapping with the nuclear envelope, defined by LAMINB1 staining (dotted blue line). Scale bar: 5 μm. **(B)** Quantitative model of HIV-1 post entry steps in activated and resting CD4 T cells. Based on previous literature, frequencies of viral particles that sequentially undergo the different post-entry steps are plotted (cell entry set arbitrarily at 100%). Data from activated CD4 T cells are fitted to a linear equation and a non-linear second order polynomial quadratic fit is displayed for data from resting CD4 T cells. **(C)** Experimental workflow for assessing early post-entry replication steps in CD4 T cells. Activated and resting CD4 T cells from autologous donors were infected with 0.5 infectious units of HIV-1 NLENG GFP p6mod reporter virus (with or without Vpx). Cells were harvested at 6, 24, 48, and 72 h post-infection (hpi) for assessment of viral nuclear import by microscopy, and viral gene expression was quantified by GFP expression at 72 hpi using flow cytometry. **(D)** HIV-1 infection quantified as the percentage of GFP+ cells at 72 hpi. Each matched pair represents a single donor. **(E)** Representative single Z-plane confocal microscopy images of infected activated and resting CD4 T cells at different time points post infection. Cells were immunostained with an anti-capsid (CA) antibody (magenta). Intranucleoplasmic capsids are indicated by arrows and are located within the nuclear boundary, defined by LAMINB1 highlighted as dotted line. Scale bar: 10 μm. **(F)** HIV-1 nuclear import, depicted as the percentage of cells bearing at least one intranucleoplasmic capsid. Images were analyzed in 3D encompassing the entire nucleus. At least 50 cells per condition per donor (n=3) were analyzed **(G)** Frequency of CA+ foci per nucleus in activated and resting CD4 T cells at 24, 48, and 72 hpi quantified in 3D.. All statistical analysis were performed as t tests comparing activated and resting conditions. Paired analysis was performed for panel D, unpaired analysis was performed for panels F and G. ns: not significant.

Post-entry barriers to productive HIV-1 infection exist at several steps of the viral life cycle in both resting and activated CD4 T cells, and incoming viral particles are progressively eliminated or inactivated by fitness constraints and cellular restriction factors. This pattern of sequential attrition prevents most particles that initially enter the cell from completing the infectious cycle. Based on quantitative analyses (Supplementary table 1), this attrition follows an approximately linear trajectory across early post-entry replication steps in activated CD4 T cells (Fig 1B, Suppl. Table 1). In resting CD4 T cells, however, additional stage-specific restrictions, most notably at reverse transcription by SAMHD1, convert this trajectory into a non-linear mode of infection (Fig 1B). T cell activation thus governs the efficiency of the HIV-1 post-entry phase.

To systematically assess this, we infected resting CD4 T cells (treated with low levels of IL-7/15 to facilitate survival without increasing permissivity to HIV-1 infection (Ananth et al. 2024) and autologous activated CD4 T cells (treated with IL-2) with 0.5 infectious units of the HIV-1 NLENG GFP reporter virus (Ananth et al. 2024). GFP expression indicates viral gene expression mainly from integrated proviruses and thus successful completion of early replication steps up to viral protein synthesis. Resting CD4 T cell infection was carried out with virus incorporating Vpx to overcome the SAMHD1 block to reverse transcription. We have previously shown that this does not interfere with nuclear import efficacy (Ananth et al. 2024) and allows investigation of all post-entry steps. Infected cells were analyzed for intranucleoplasmic capsids (CA) at 6, 24, 48, and 72 hours post-infection (hpi) by microscopy, and infection rates were measured by GFP expression using flow cytometry at 72 hpi (Fig 1C). Although the cells were infected with similar infectious units as titrated on TZM-bl reporter cells, infection rates were moderately, but significantly lower in resting CD4 T cells at 72 hpi (Fig 1D). The lower infection rates in resting CD4 T cells likely reflect the barrier imposed by the limited remodeling of the cortical actin network despite spinoculation, which restricts viral entry (Yoder et al. 2008). Infected activated and resting CD4 T cells were immunostained with an anti-CA antibody (Muller et al. 2021) to visualize intact or broken HIV-1 capsid structures in the nucleoplasm (Fig. 1E). HIV-1 nuclear import efficiency was quantified as the percentage of cells showing at least one intranucleoplasmic HIV-1 CA focus. Nuclear import of CA was not observed up to 6 hpi (Fig 1F) but was robust at later time points (24 to 72 hpi) in both resting and activated CD4 T cells (Fig 1E, F). In line with our previous study (Ananth et al. 2024), this confirms that nuclear import of HIV-1 capsids in resting CD4 T cells is not restricted following cell-free infection. Barriers to HIV-1 nuclear import in resting CD4 T cells, as recently reported for HIV-1 infection following priming of CD4 signaling (Mesner et al. 2026), and the effect of CD4 priming on viral nuclear import and infection were not observed in our experimental system. This may reflect differences in culture conditions, including the nature and quantity of added cytokines such as IL-2, influencing the permissivity of phenotypically resting CD4 T cells for HIV-1 infection. In our experiments, nuclear import efficiency was significantly higher in resting CD4 T cells at 24 and 48 hpi, and the frequency of cells with more than one intranuclear CA focus was elevated in resting over activated CD4 T cells despite the lower infection rates prior to T cell activation (Fig 1F, G). These differences became less pronounced at 72 hpi. Vpx did not influence capsid import in resting CD4 T cells, as spinoculation with identical amounts of integration-deficient HIV-1 (NNHIV) in the absence of Vpx also showed significantly higher nuclear import in resting compared to activated CD4 T cells at 24 hpi (Suppl. Fig. 1B). These results demonstrate that HIV-1 CA nuclear import is not restricted in resting CD4 T cells but rather efficient compared to activated cells.

### NUP62 acts as a T cell activation-dependent molecular switch for HIV-1 infection

HIV-1 nuclear import is a well-orchestrated process that involves sequential interaction of the capsid with host cell proteins, including components of the NPC, most importantly NUP358 on the cytoplasmic tail; NUP35, NUP62, and NUP98 assembled in the central pore; and TPR and NUP153 in the nuclear basket (Fig 2A) (Zila et al. 2021). To further examine the relationship between viral nuclear import and infection upon T cell activation, we investigated whether HIV capsid import in resting or activated CD4 T cells requires specific NPC components. Comparing bulk NUP protein levels revealed significant variation in individual NUP abundance between the two activation states, and the magnitude of these differences varied between cells from different donors tested. While NUP54 and NUP358 were largely unaffected by T cell activation, levels of the established HIV-1 nuclear import factors NUP62 and NUP153 were substantially increased by CD4 T cell activation, and TPR and NUP35 were more abundant in resting CD4 T cells (Suppl. Fig 1C, D). T cell activation thus differentially impacts expression and/or stability of individual NPC components. To investigate the impact of T cell activation on the role of these factors in HIV-1 post-entry steps, we targeted individual *NUP* genes by activation-neutral gene editing in resting CD4 T cells (Albanese et al. 2022) (Supplementary table 2) and incubated them until the respective protein was depleted, as determined by Western Blot analysis (Fig 2B). While knock out (KO) of *NUP62*, *NUP98*, *NUP153* or *TPR* led to efficient protein depletion relative to control cells treated with non-targeting (NT) guides in cells from all donors tested 2–3 weeks post gene editing (Fig. 2C, Suppl. Fig. 2A), depletion efficiency was donor-dependent for NUP358 (Fig. 2C) and depletion of NUP54, NUP58, or NUP35 was either inefficient or non-reproducible across donors (Suppl. Fig. 2A). For responsive NUPs, the protein depletion detected by Western Blotting translated to a strong reduction of the NPC-associated protein at the single cell level (see Suppl. Fig 2B, C for *NUP62* and *NUP153* KO). For NUP62, transcriptome analysis documented high KO efficacy at the genomic level (Suppl. Fig. 2D). Depletion of some NUPs, including NUP98, which is present at the highest copy number per NPC, strongly reduced cell viability, whereas depletion of most other NUPs had only moderate or no cytotoxic effect (Fig 2D). Based on reliable KO efficiency and cell viability, further infection and microscopy analysis were performed for NUP62, NUP153 and NUP358. To this end, resting CD4 T cells with validated protein depletion were either activated or kept resting for 72 h, infected, and functionally characterized 48 or 72 hpi (Fig 2E). KO of *NUP62* strongly reduced viral infection levels in activated CD4 T cells (mean difference of 7.5-fold) compared to NT control cells. In contrast, infection rates were significantly higher (mean difference of 2.1-fold and a maximum of 2.5-fold in one donor) in resting CD4 T cells in the absence of NUP62 compared to control (Fig 2F). Thus, NUP62 depletion can increase the permissivity of resting CD4 T cells relative to activated CD4 T cells by up to 18-fold. Importantly, this increased permissivity of resting CD4 T cells for HIV-1 infection was not associated with detectable cell activation as evaluated by cell surface activation marker display or gene expression profile (Suppl. Fig 3A-C). This effect was specific to NUP62 since KO of known nuclear import factors NUP358 and NUP153 markedly reduced HIV-1 infection of activated CD4 T cells but did not increase infection levels (moderate reduction, not statistically significant) of resting CD4 T cells (Suppl. Fig 3D, E). The differential effect of HIV-1 infection in *NUP62* KO cells was independent of Vpx as the presence of Vpx in activated CD4 T cells did not abrogate the requirement of NUP62 for infection and NUP62 depletion was not sufficient to bypass the SAMHD1 barrier in the absence of Vpx in resting CD4 T cells (Suppl. Fig. 3F). To test if the ability of activated CD4 T cells to proliferate is critical for the effect of NUP62 on HIV infection, we generated *NUP62* KO in monocytes and terminally differentiated them into non-proliferative macrophages and Langerhans cells (Fig 2G). Following verification of target protein depletion by western blot (Suppl. Fig. 3G), the cells were infected with a R5-tropic HIV-1 reporter virus (Fig 2G). Similar to proliferating activated CD4 T cells, NUP62 depletion markedly reduced HIV-1 infection in both cell types (Fig 2H, I), suggesting T cell activation parameters other than induction of cell proliferation as regulators of NUP62 function. These results identify NUP62 as a molecular switch that regulates HIV-1 infection in a T cell activation-dependent manner.

**Figure 2.**
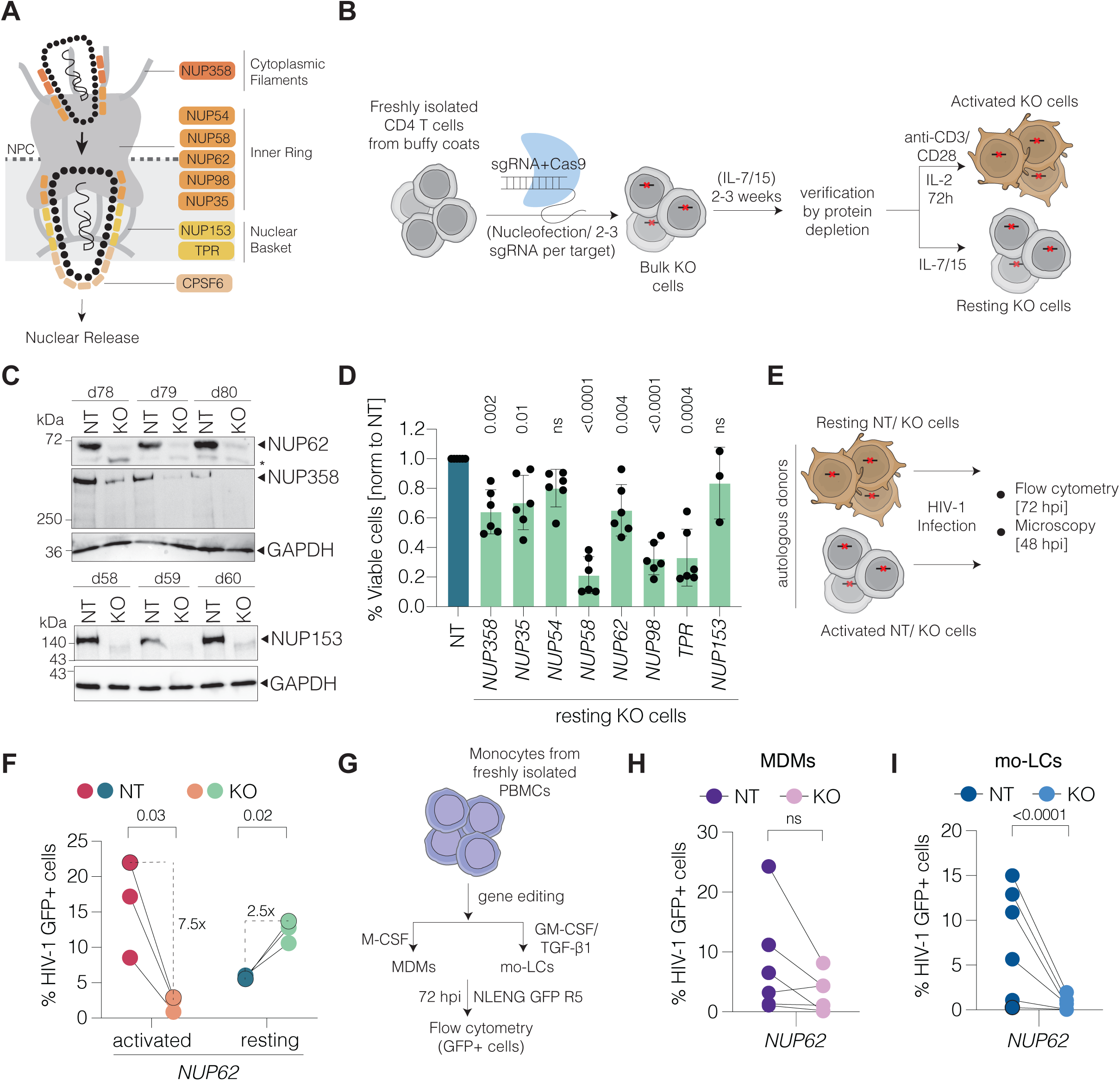
T cell activation switches NUP62 from a dependency to inhibitory factor for HIV-1 infection. **(A)** Schematic representation of HIV-1 traversing the nuclear pore complex (NPC) via interactions with nucleoporins (NUPs). NUPs forming the scaffold (cytoplasmic filaments and nuclear basket) and the inner ring (FG-NUPs) are highlighted. **(B)** Schematic workflow for gene editing in primary resting CD4 T cells. Freshly isolated CD4 T cells from buffy coats were nucleofected with multiple sgRNAs targeting a single gene in complex with Cas9. After 2–3 weeks, successful KO in bulk cells was verified by Western blot. A fraction of these cells was activated with anti-CD3/CD28 Dynabeads for 72 h to generate a pool of activated CD4 T cells autologous to the gene-edited resting CD4 T cells. **(C)** Representative western blot of gene-edited NT and KO cells for the targets *NUP62*, *NUP358* and *NUP153* from three independent donors. **(D)** Viability of gene-edited resting CD4 T cells after ∼2.5 weeks measured by flow cytometry using viability dye. Values were normalized to NT, which was arbitrarily set to 1. **(E)** Activated and resting non-targeting (NT) control and NUP-targeted knockout (KO) CD4 T cells were infected with the HIV-1 NLENG GFP reporter virus lacking Vpx (activated) or containing Vpx (resting). Flow cytometry and microscopy analysis were performed at 72 and 48 hpi. **(F)** Infection rates, measured as the percentage of GFP-expressing cells in NT and *NUP62* KO activated and resting CD4 T cells. Data shown here are for three independent donors. **(G)** Schematic workflow for the generation of gene-edited monocyte-derived macrophages (MDMs) and monocyte-derived Langerhans cells (mo-LCs). Monocytes were isolated from PBMCs and nucleofected with sgRNA–Cas9 complexes targeting *NUP62*. Following nucleofection, cells were seeded in 96-well plates and differentiated for 7 days into MDMs or mo-LCs using macrophage colony-stimulating factor (M-CSF) or granulocyte– macrophage colony-stimulating factor (GM-CSF), respectively. Differentiated cells were then infected with an R5-tropic NLENG reporter virus, and infection was quantified by GFP expression at 3 days post-infection. **(H, I)** Infection rates at 72 hpi in gene-edited monocyte-derived macrophages (MDMs) (H) and monocyte-derived Langerhans cells (mo-LCs) (I). Statistical significance was calculated using one sample t test for panel D and paired t test for panels F, H and I. ns: not significant.

### NUP62 alters early post-entry steps in resting CD4 T cells

We next examined the effect of *NUP62* KO on HIV-1 nuclear import by quantifying nuclear capsid localization. In activated CD4 T cells, *NUP62* KO slightly reduced the number of cells with nuclear capsid foci in cells from some donors, but this effect was overall not significant (Fig 3A, B). However, the number of capsid foci per nucleus remained significantly lower upon *NUP62* KO in these cells (Fig. 3C). In contrast, *NUP62* KO significantly increased the proportion of resting CD4 T cells containing intranucleoplasmic capsid foci as well as the number of capsid foci per nucleus (Fig. 3A-C). Enhanced viral nuclear import in NUP62-depleted resting CD4 T cells was not associated with global changes in the spatial distribution of NPCs as detected by super-resolution microscopy (Suppl. Fig. 4A–C) or altered barrier integrity as assessed by fluorescent dextran permeability assays (Suppl. Fig. 4D-F). However, NUP62 depletion differentially altered the overall abundance of other NUPs that are found in complex with NUP62 in the inner pore and cytoplasmic regions of the NPC in activated and resting CD4 T cells (Suppl. Fig. 4G–I). NUP62 depletion thus likely adds to the differences in NPC composition and heterogeneity between activated and resting CD4 T cells. We conclude that the effects of NUP62 depletion on HIV-1 nuclear import and infection do not result from gross alterations in NPC distribution or barrier integrity but may reflect changes in NPC composition.

**Figure 3.**
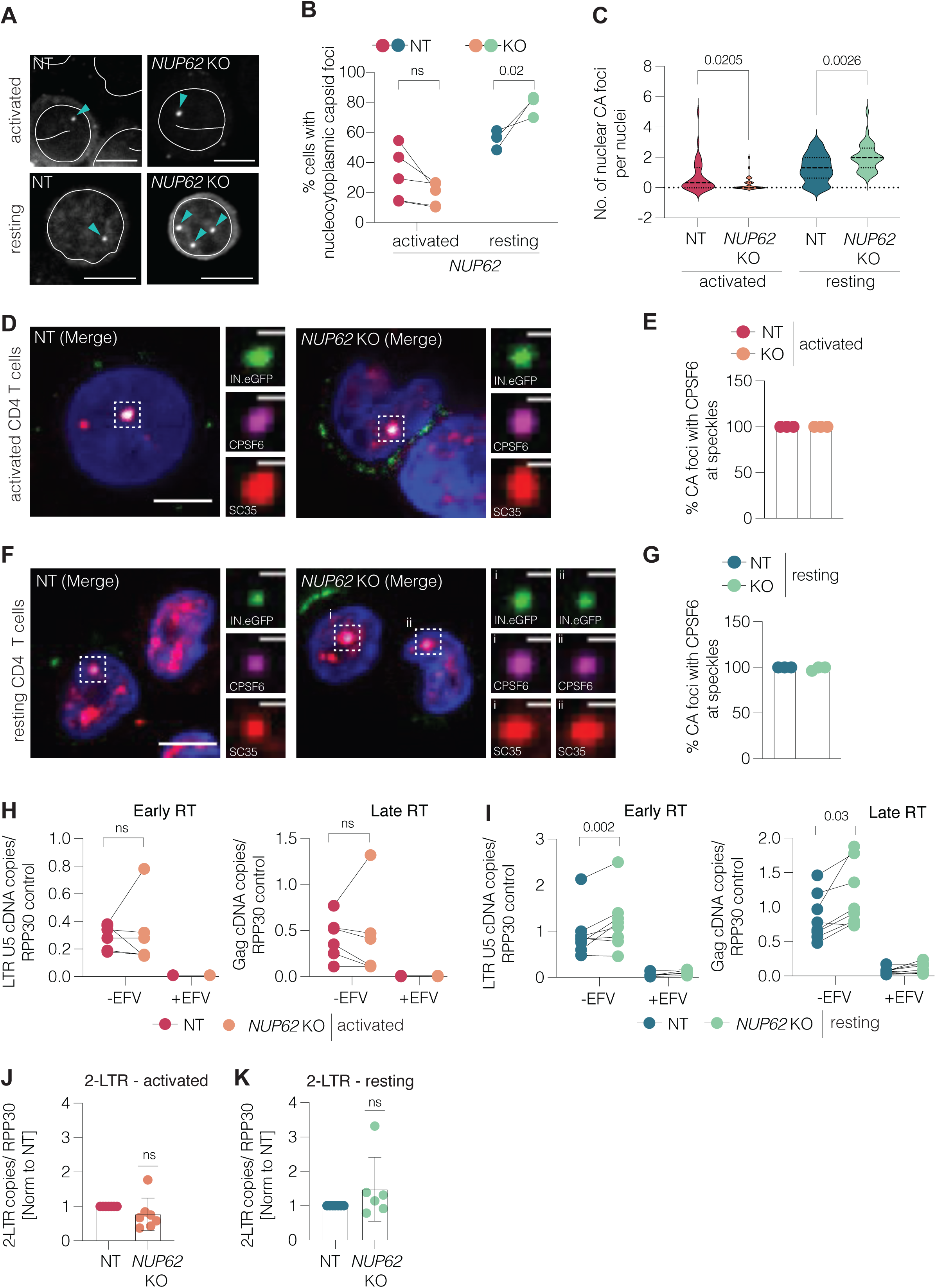
Nuclear import and reverse transcription are boosted upon *NUP62* KO in resting CD4 T cells. **(A)** Representative single Z-slice confocal images of NT and *NUP62* KO activated and resting CD4 T cells showing intranucleoplasmic capsids (CA highlighted by cyan arrow, LAMINB1 boundary by white line). Scale bar: 5 μm. **(B)** Nuclear import efficiency quantified as % NT and *NUP62* KO cells with at least one capsid positive focus. **(C)** Quantification of CA events per nucleus in NT and *NUP62* KO activated and resting CD4 T cells. **(D–G)** Nuclear localization of viral capsids in activated and resting CD4 T cells. (D, F) Representative single Z-slice confocal images of activated and resting NT and *NUP62* KO cells infected with HIV-1 NL4.3 WT (IN.eGFP) virus and immunostained for CPSF6 (magenta) and SC35 (Cornish et al.) as a marker for nuclear speckles (Cornish et al.). The nucleus was stained with Hoechst (blue). Zoomed images focus on intranucleoplasmic HIV-1 CA events highlighted by the white box. Scale bar: 5 μm; zoom scale bar: 1 μm. (E, G) Frequency of HIV-1 CA colocalization with both CPSF6 and SC35 in activated and resting NT and *NUP62* KO CD4 T cells. **(H, I)** Quantification of HIV-1 reverse-transcribed products by digital droplet PCR (ddPCR) at 72 hpi in activated (H) and resting (I) CD4 T cells. Early and late reverse transcription (RT) products were defined by the number of LTR U5 and Gag cDNA copies, respectively. Cells treated with Efavirenz (EFV) were used as a control. n ≥ 6 independent donors. **(J, K)** 2-LTR circles quantification in NT and *NUP62* KO activated and resting CD4 T cells by ddPCR. Statistical significance was determined using a ratio-paired t test for panels B, H and I, and one sample t-test for panels J and K. ns: not significant.

Following nuclear entry, HIV-1 capsids retained the ability to recruit CPSF6 and localize to nuclear speckles with the same frequency in the absence or presence of NUP62 in both activated and resting CD4 T cells, as assessed by immunofluorescence microscopy (Fig. 3D–G). Therefore, the trajectory of the virus from the NPC to subnuclear speckles and the accessibility of capsid for interaction with CPSF6 are independent of NUP62. Furthermore, capsids carrying the A77V or N74A mutation were impaired in capsid release into the nucleoplasm and nuclear entry, respectively (Suppl. Fig. 5A). CA foci were largely found at the nuclear envelope and exhibited markedly reduced nuclear import irrespective of NUP62 depletion (Suppl. Fig. 5B–E). NUP62 depletion therefore does not bypass the viral determinants required for nuclear entry and subsequent release into the nucleoplasm.

Recent studies revealed that viral reverse transcription and nuclear import are temporally overlapping processes, with reverse transcription initiating in the cytoplasm and completing in the nucleus following nuclear entry (Dharan et al. 2020, Francis et al. 2020). Consistent with the moderate effects on capsid import, quantifying early and late reverse transcription products revealed that NUP62 depletion did not alter the abundance of either product in activated CD4 T cells (Fig 3H). In contrast, both early and late viral cDNA levels were increased in resting cells in the absence of NUP62 (Fig 3I), matching the increase in viral nuclear import. These results suggest that the enhanced nuclear import in resting cells lacking NUP62 boosts detectable reverse-transcribed intermediates and that early reverse transcription may occur after nuclear import in resting CD4 T cells. While reverse transcription initiates in the cytoplasm and is completed in the nucleus in most HIV-1 target cells (Dharan et al. 2020, Francis et al. 2020, Selyutina et al. 2020), features of resting CD4 T cells such as reduced nuclear-cytoplasmic ratio (Fig 1A) and incomplete restoration of dNTP pools even in the presence of Vpx (Baldauf et al. 2012) may alter dynamics of reverse transcription with respect to nuclear import. Quantification of 2-LTR circles, an indicator of complete reverse transcribed products in the nucleus, revealed no significant difference upon *NUP62* KO both in activated and resting CD4 T cells (Fig 3J, K). Therefore, the moderate increase in reverse transcription products in resting CD4 T cells did not affect the overall availability of complete vDNA for subsequent infection stages. Together, these results establish NUP62 as a regulator of early post-entry steps that is dispensable for the availability of completed reverse transcription products in resting CD4 T cells.

### NUP62 influences selective integration of proviruses in resting CD4 T cells

We next conducted integration-site sequencing to evaluate quantitative and qualitative impacts of NUP62 depletion on HIV-1 integration. The number of unique HIV-1 integration sites recovered by IS sequencing did not differ significantly between NT and *NUP62* KO CD4 T cells irrespective of their activation state, suggesting that the similar amounts of vDNA produced were comparably competent for viral integration (Fig. 4A, B). We next examined whether NUP62 depletion affected the genomic distribution HIV-1 integration sites. Spatial position inference of the nuclear (SPIN) state analysis was performed to classify HIV-1 integration sites (ISs) based on the different nuclear compartments (Gholamalamdari et al. 2025). Expectedly, HIV-1 integrations were enriched within the high gene-dense speckle-associated domains in all samples analyzed and generally depleted from the lamina state (28.2% of IS vs 8.1% of random matched control sites in activated NT cells; lamina 9.7% vs 23.7%) (Fig 4C, Suppl. Table 3) (Marini et al. 2015, Sowd et al. 2016, Achuthan et al. 2018, Bedwell et al. 2021). However, ISs showed a slight preference for integration into lamin and lamin-associated regions in resting NT cells over activated NT cells (40.7% vs 32.7% of ISs) and this redistribution was significant (p=1.3×10^−5^, Fig 4C, Suppl. Fig. 6A). *NUP62* KO significantly altered this distribution in resting (p = 4.3 × 10⁻⁹) but not in activated CD4 T cells (p=0.23) (Fig 4C, Suppl. Fig. 6A). In resting cells, NUP62 depletion redistributed ISs from speckle-associated and interior-active states (Speckle 21.0% to 18.8%; Interior_Act3 10.6% to 9.2%) toward lamina-proximal states (Near_Lm2 15.3% to 17.4%, Lamina 14.5% to 16.7%; all p<0.01) (Suppl. Fig. 6A), increasing the fraction of ISs outside speckle-associated domains from 79.0% to 81.2%. The interior repressive states were unaffected, indicating a redistribution toward the nuclear periphery rather than a shift into repressed chromatin at comparable radial position.

**Figure 4.**
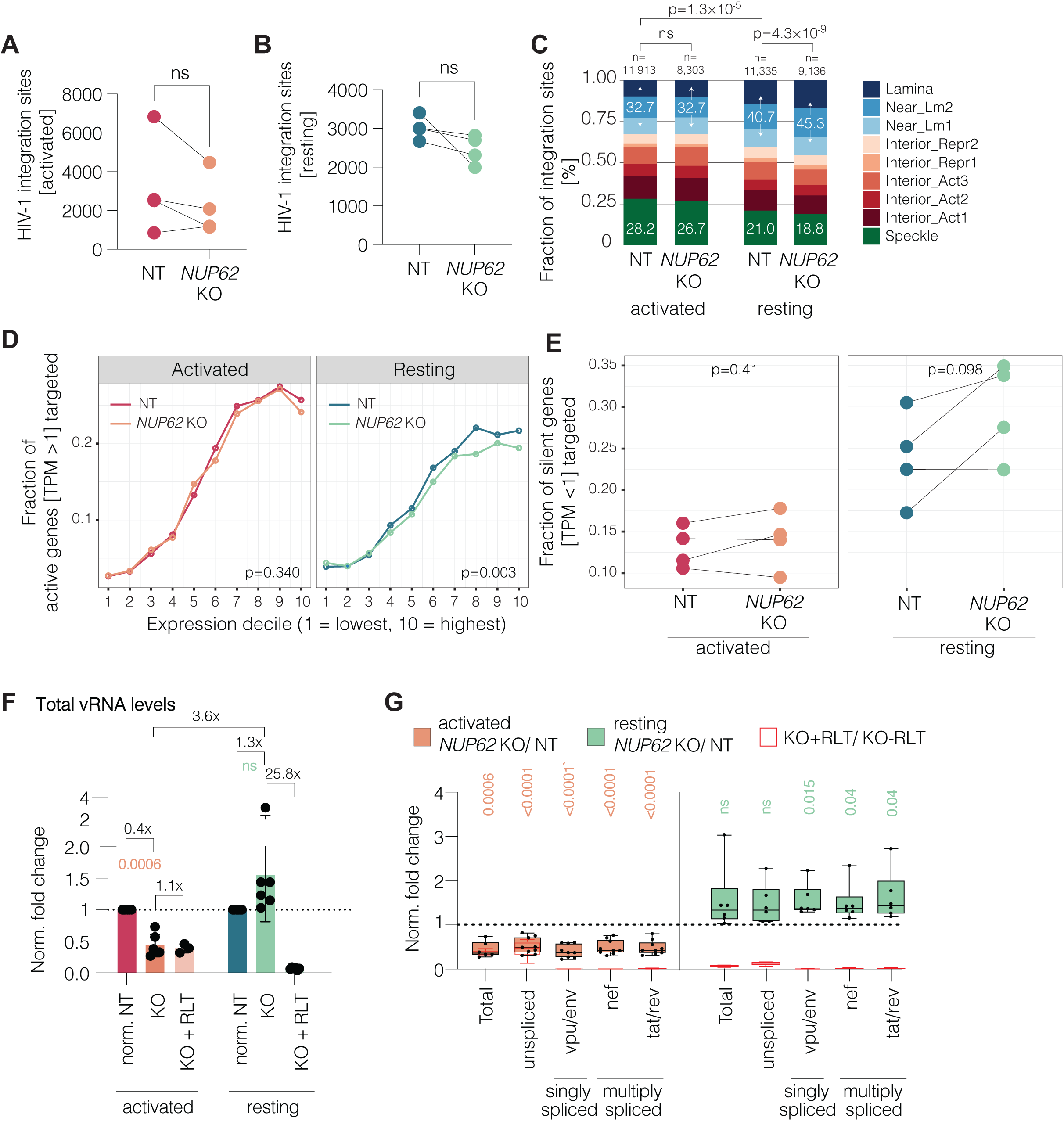
NUP62 controls the genomic positioning of HIV-1 and viral transcription in CD4 T cells. (A, **B)** Total HIV-1 integration sites (ISs) from integration-site sequencing in NT and *NUP62* KO activated and resting CD4 T cells at 72 hpi. **(C)** Spatial Position Inference of the Nuclear genome (SPIN) state analysis of HIV-1 ISs correlating the frequency of integration with distinct nuclear compartments classified based on active or gene-dense repressive chromatin regions. P-values calculated using χ² tests of independence across the nine states **(D)** Fraction of active genes targeted by ISs after equalizing for number of integrations. All active genes (TPM > 1) identified from RNA-seq data of infected NT and *NUP62* KO activated and resting CD4 T cells were split into 10 bins. 1 = least active; 10 = most active. P values were calculated based on generalized additive model (GAM) curves. **(E)** Fraction of silent genes (TPM < 1) targeted by HIV-1 ISs in NT and *NUP62* KO activated and resting CD4 T cells. **(F)** Quantification of total HIV-1 RNA from NT and *NUP62* KO activated and resting CD4 T cells, represented as fold change over NT for KO samples (NT set to 1), and fold change over KO for KO cells treated with Raltegravir (KO set to 1). Statistical significance was calculated by comparing NT vs KO (highlighted in different colors for resting and activated conditions) and fold difference in the effect is depicted **(G)** Quantification similar to panel F for different unspliced and spliced viral transcripts. Significance was calculated for the fold change between KO and NT using a one-sample t-test where NT was arbitrarily set to 1. Values of RLT-treated KO samples were calculated as in panel F but represented in red. Statistical analysis was calculated using paired t test for panel A, B and E.

Since this analysis cannot resolve the contribution of a small set of ISs towards viral and/or cellular transcription, we specifically mapped IS locations to transcriptionally active and silent genic regions using the cellular gene expression profile of infected NT and *NUP62* KO activated and resting CD4 T cells determined by RNAseq. Transcripts of *egfp* and *env*, expressed from the viral promoter, showed a trend toward decreased expression in *NUP62* KO activated CD4 T cells and increased expression in *NUP62* KO resting CD4 T cells (Suppl. Figs. 6B, C). These differences were moderate compared to the effect of NUP62 on viral gene expression (Fig 2F) suggesting that post-transcriptional processes, RNA stability, and export may add to the role of NUP62 in infection. Cellular genes with transcripts per million [TPM] >1 were classified as active, and TPM <1 as silent. Consistent with the known preference of HIV-1 for active chromatin (Brady et al. 2009) (Lucic et al. 2019), the fraction of integrations increased with gene expression level in active genic regions across expression deciles (1 = least active; 10 = most active) in both resting and activated cells (Fig. 4D). Importantly, NUP62 depletion in resting CD4 T cells significantly reduced the fraction of integrations into highly active genes (upper expression deciles) (Fig. 4D). In contrast, the distribution of integrations in active genes was indistinguishable between activated NT control and *NUP62* KO CD4 T cells. This suggests that depletion of NUP62 redirects HIV-1 integration away from highly active genes specifically in resting CD4 T cells. Accordingly, the fraction of integration sites within silent genes was higher in *NUP62* KO than in NT resting CD4 T cells in three of the four donors tested, albeit not significant (Fig. 4E), and such differences were not observed in activated CD4 T cells. Together, these results indicate that NUP62 can contribute to the spatial positioning of the provirus specifically in resting CD4 T cells by redirecting a subset toward silent genic regions.

### NUP62 depletion affects levels of HIV-1 transcripts in a T cell activation-dependent manner

We next assessed the effect of NUP62 depletion on viral transcript abundance. KO of NUP62 only altered the expression of few cellular genes with no global changes in the transcriptome (Suppl. Fig. 6D, E). However, *NUP62* KO in activated CD4 T cells significantly reduced total HIV-1 mRNA to 40% of that of NT control cells. This inhibition was similar to that observed in *NUP62* KO activated CD4 T cells treated with raltegravir (RLT) during infection (Fig. 4F). Since GFP expression was completely abrogated in these cells (Suppl. Fig. 6F), the detection of residual viral transcripts in RLT-treated activated CD4 T cells can be attributed to viral genomes from input virus rather than incomplete inhibition of infection. In resting CD4 T cells in contrast, *NUP62* KO increased total viral mRNA levels to an average of 130% of that in NT control cells and up to 25-fold over the RLT control in cells from all donors tested. The magnitude of this effect was subject to donor-to-donor variability but collectively, total vRNA transcripts were on average 3.6-fold enriched in resting over activated CD4 T cells when NUP62 was depleted (Fig. 4F). Changes in viral splice-variant abundance paralleled those in total HIV-1 mRNA in both activation states (Fig. 4G), suggesting that NUP62 depletion does not substantially alter viral mRNA splicing. Together, these findings establish the control of viral transcript abundance as an unexpected additional mechanism by which NUP62 exerts T cell activation-dependent regulation of the HIV-1 life cycle.

## Discussion

This study aimed to provide insight into the mechanism that allows efficient HIV-1 nuclear import in resting CD4 T cells despite their low number of NPCs and closed chromatin architecture and its impact on viral replication. Applying our recently developed activation-neutral gene editing method enabled us to efficiently deplete individual NUPs in resting CD4 T cells and to identify three NUPs that can be depleted without marked effects on cell viability. The study focused on NUP62, which could be depleted without compromising NPC distribution and permissivity, or T cell activation state. Comparing resting and activated CD4 T cells from the same donor revealed an unexpected dichotomy in the role of NUP62 in HIV infection. NUP62 facilitates HIV-1 infection in activated CD4 T cells as measured by synthesis of new viral proteins. In contrast, NUP62 presents a barrier to HIV infection in resting CD4 T cells. This finding reconciles previous reports from cell line models that NUP62, in addition to its prototypic role as an HIV dependency factor, can also limit HIV infection in e.g HT1080 epithelial cells (Kane et al. 2018, Xue et al. 2023) and establishes that a physiological stimulus can switch between the pro- and antiviral roles of NUP62 in HIV-1 infection of primary human target cells. Since our results in non-proliferating myeloid cells exclude cell proliferation as the driver of this functional switch, this regulation is likely a specific consequence of T cell activation. Mechanistically, it was surprising that NUP62 depletion only exerts moderate effects on nuclear import efficiency of capsids, which does not translate to changes in subsequent steps such as nuclear localization of the subviral complex and/ or increased efficacy of reverse transcription completion. Instead, NUP62 affects steps further downstream in the HIV life cycle such as IS selection and abundance of viral transcripts in an activation state-dependent manner. These results reveal that NUP62 exerts important functions subsequent to nuclear entry of capsids and that this post-nuclear import role of NUP62 is regulated by T cell activation.

Our study reveals a dual role of NUP62 in HIV-1 replication in primary human CD4 T cells that includes direct effects during nuclear import but also on downstream steps in the viral life cycle. Effects on nuclear import efficiency were surprisingly moderate, which is consistent with our finding that NPC density remains unaltered in the absence of NUP62. More subtle ultrastructural changes to NPC architecture and NUP stoichiometry likely explain the quantitative changes at the level of HIV-1 nuclear import in resting CD4 T cells. The stoichiometry of NUP62 at the NPC and NPC elasticity dictate the extent of capsid penetration into the nucleus (Dharan et al. 2020, Shen et al. 2023, Hou et al. 2025, Kreysing et al. 2025). Our observation that not all NUPs are upregulated upon T cell activation despite the concomitant increase in NPC copy number suggests that the stoichiometric composition of NUPs within the NPC can differ between resting and activated CD4 T cells. Mechanistically distinct pathways that govern NPC assembly during cell proliferation and dynamics of assembled NPCs might contribute to these differences (Rabut et al. 2004, Otsuka et al. 2023). NPC-free NUP62 that interacts with cytoplasmic subviral complexes (Monette et al. 2011, Dharan et al. 2020) and virus-induced remodeling of NPC composition (Porter and Palmenberg 2009, Xue et al. 2024, Mesner et al. 2026) may add regulatory layers to the divergence of the NUP62 function in nuclear import at different cell activation states.

In addition, NUP62 may also act as functional capsid selectivity barrier that provides capsids with an experience during nuclear import that affects downstream replication steps. This may include that the lack of NUP62 alters mechanical properties such as NPC elasticity (Hou et al. 2025) (Kreysing et al. 2025). However, the effect of NUP62 on these late replication stages may go beyond its role at the NPC. NUP62 may directly affect post-nuclear entry replication steps and/or indirectly alter them through involvement of cellular cargo whose nucleo-cytoplasmic distribution is regulated by the activation-induced switch in NUP62 function. Testing the involvement of NUP62-dependent cargo molecules will require unbiased proteomics approaches to dissect the impact of NUP62 on nuclear-cytoplasmic protein distribution in our experimental system. In addition, import-independent roles of NUP62 such as the regulation of chromatin organization (Breuer and Ohkura 2015) and/ or cellular RNA export (Dargemont et al. 1995, Terry and Wente 2007, Monette et al. 2011) may directly affect post nuclear import replication steps and the impact of NUP62 on viral RNA transcription, stability, export and/or protein synthesis warrants further investigation. Future studies will aim at dissecting the contribution of these not mutually exclusive mechanisms to the regulation of HIV-1 infection of CD4 T cells by NUP62.

The finding that, in cells from the majority of donors tested, the increase in viral transcript abundance and gene expression upon NUP62 depletion is associated with a shift of a small subset of proviral integrations (4.6% of ISs relocating out of interior-active states) to silent genic regions in resting CD4 T cells is counterintuitive. The SPIN state analysis shows that NUP62 depletion shifted ISs toward lamina-proximal states while leaving the interior repressive states unchanged, indicating a redistribution toward the nuclear periphery rather than into interior repressive chromatin. Speckle-associated targeting was reduced but not abolished, arguing that NUP62 repositions proviral integrations. Together, these findings raise the possibility that in resting CD4 T cells, NUP62 influences the positioning of the provirus to prioritize transcriptional competence of ISs despite the generally low accessibility of chromatin. Such rewiring of IS selection could potentially be beneficial for the virus. First, it avoids competition for cellular resources such as transcription factors at sites of high transcriptional activity, which may be limiting in resting CD4 T cells that are not primed for increased cellular function. Second, preselection of specific sites, for example those that may retain small nanoclusters of active RNA Pol II even amid globally low transcriptional activity (Xu et al. 2024), may sustain efficient transcription irrespective of the surrounding gene expression landscape. This model is consistent with evidence that transcriptionally silent latent HIV-1 proviruses maintain a primed state through continuous replacement of paused RNA Pol II, enabling rapid transcriptional induction upon activation (Jadlowsky et al. 2014). Robust viral transcription may therefore not be strictly confined to actively transcribing loci. Loss of NUP62, possibly by disrupting interactions of chromatin-bound NUP62 with viral integrase (Ao et al. 2012), may reroute integration-competent viral cDNA toward transcriptionally silent, yet Pol II-accessible chromatin regions in resting CD4 T cells. Since only a fraction of proviruses contribute to the generation of viral transcripts (Einkauf et al. 2022), the shift of a small subset of integrations in such an environment that is generally repressive for host cell transcription but rendered permissive for the expression of viral genes may account for the observed increase in viral transcripts in resting CD4 T cells.

Our results also reveal that the efficiencies of nuclear import, reverse transcription and amounts of viral transcripts are not directly proportional to one another. This was particularly apparent in resting CD4 T cells, where NUP62 depletion increases nuclear import with a concomitant enhancement of reverse transcribed intermediates, but this is not converted into improved efficiency of reverse transcription completion or integration. Yet, NUP62 depletion elevates viral transcript abundance in resting CD4 T cells. Summarizing these results in the form of regression model curves revealed that NUP62-dependent effects upon T cell activation deviate HIV-1 replication steps from apparent linear attrition (Fig 1B) to a non-linear model of infection (Fig 5A). Replication kinetics upon NUP62 depletion in CD4 T cells are better described by an ‘hourglass model’ of infection where consistent differences of varying magnitudes are observed in early and late steps even though intermediate stages remain unaffected. Here, early post-entry as well as pre- and post-integration steps are quantitatively altered to varying magnitudes upon T cell activation and are regulated by NUP62 (Fig 5A, B). This is in line with recent findings that NPC heterogeneity has functional implications for nuclear import of HIV capsids and can uncouple the efficacy of nuclear import from that of viral gene expression (Dharan and Campbell, personal communication). Similar observations were made for the HIV-1 nuclear import cofactor CPSF6: While CPSF6 is required for early post-entry steps that include capsid release into the nucleus and thus indispensable for viral nuclear import (Achuthan et al. 2018, Bejarano et al. 2019), lack of CPSF6 boosts HIV-1 infection downstream due to reduced expression of innate immune factors as a result of defective polyadenylation function in activated CD4 T cells (Cornish et al. 2025). In resting CD4 T cells however, CPSF6 is indispensable for infection (Albanese et al. 2022), suggesting that T cell activation regulates how CPSF6 governs HIV-1 infection. Since CPSF6 depletion has the opposite effect on HIV-1 infection of CD4 T cells than depletion of NUP62 and the infectivity of CPSF6-binding deficient capsids cannot be rescued by NUP62 depletion in resting CD4 T cells, the effects of CPSF6 and NUP62 are likely mechanistically distinct. This increasing number of observations of non-linear downstream consequences of alterations in nuclear import of HIV capsids supports the concept that NPC components are multifunctional and can imprint qualitative changes on HIV capsids or alter cellular functions that manifest only at later stages of the viral life cycle.

**Figure 5.**
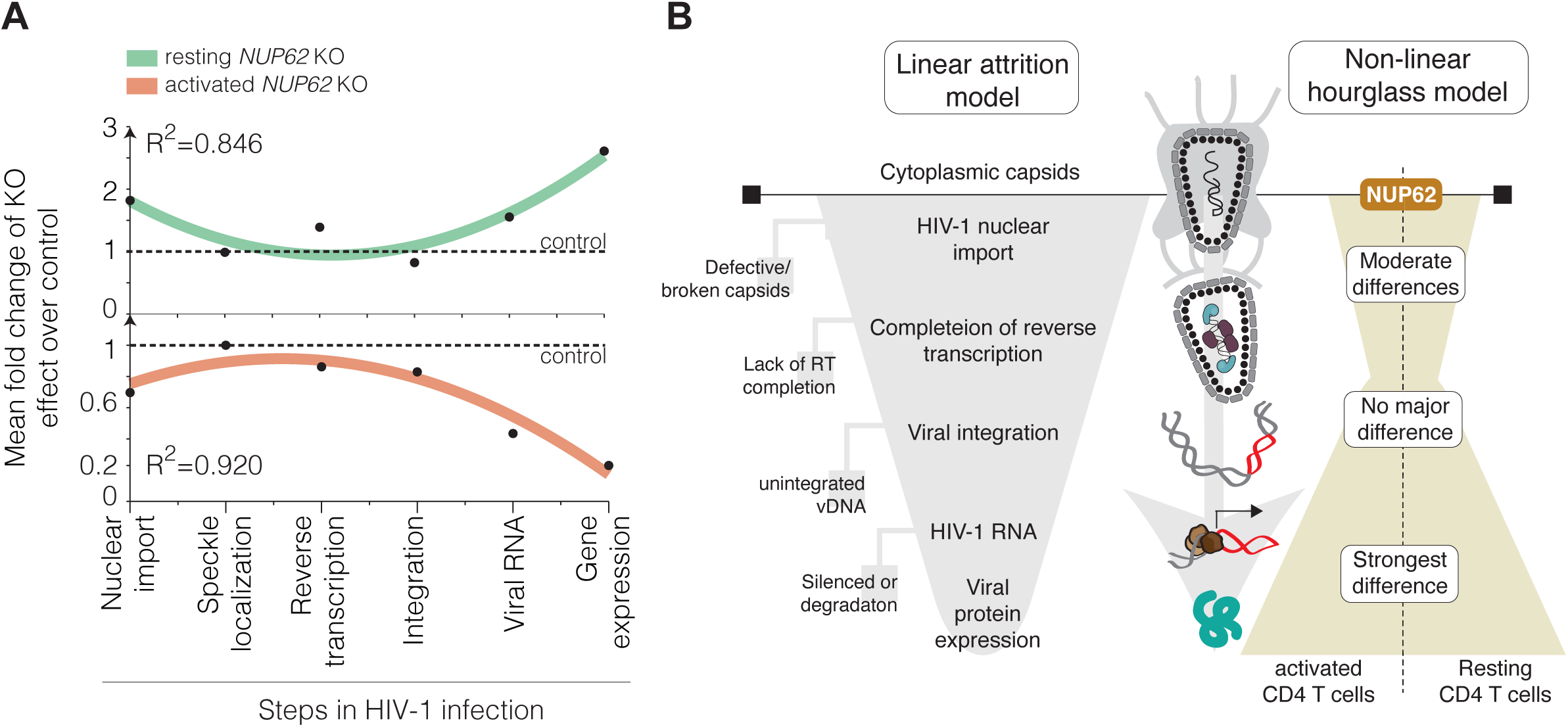
T cell activation alters the HIV-1 infection model in CD4 T cells. **(A)** Mean fold change of the *NUP62* KO effect on the different HIV-1 post-entry steps in activated and resting CD4 T cells. Non-linear relationships between the different stages were modeled using second-order polynomial regression. The root mean square deviation (R2) is shown. **(B)** Graphical summary: T cell activation switches the quantitative HIV-1 infection model from a downward funnel of restriction that follows linear attrition to a non-linear hourglass-like model using NUP62 as a molecular switch. NUP62-dependent changes to T cell activation were moderate for early post entry steps including HIV-1 nuclear import and reverse transcription. However, later replication stages such as generation of viral transcripts and gene expression were strongly altered. A qualitative impact was also seen at the level of integration in resting CD4 T cells through repositioning of the viral genome towards silent genic regions.

Together, our study establishes NUP62 as an orchestrator of post-nuclear entry steps in CD4 T cells that is regulated by T cell activation. We propose that HIV-1 exploits this functional plasticity of NUP62 to adjust strategies for provirus integration and viral gene expression to the distinct NPC abundance, composition and architecture as well as nuclear and genome organization in resting and activated CD4 T cells.

## Acknowledgments

We thank Lopamudra Sadhu for designing NUP153 sgRNAs, the IDIP imaging platform for access to advanced microscopy, and Severina Klaus for specific support. We would also like to thank Bruno Srezovic, Exaltum, LTD for their excellent bioinformatic support services. We gratefully acknowledge Carolyn Larabell and Mark Le Gros from the National Center for X-ray Tomography (NCXT), Advanced Light Source (ALS), Lawrence Berkeley National Laboratory, Berkeley for their support with soft X-ray tomography data acquisition. FA and VW are supported from the MSCA-DN CLEXM project (grant agreement No. 101120151) and ERC SyG NanoX (grant agreement No. 101167089). This project is supported by the Deutsche Forschungsgemeinschaft (DFG, German Research Foundation) under Germany’s Excellence Strategy – EXC-3018/1 – 533587280 (Cluster of Excellence SynthImmune), project 568772942 – FA 378/28-1) and project 240245660-SFB 1129, project 5 (HGK), project 6 (BM) and project 20 (ML) and the German Centre for Infection Research (DZIF) (TTU 04.820 – HIV reservoir to OTF and ML) and (TTU04.709 –Preclinical HIV Research to ML).

## Author contributions

Conceptualization: OTF; Methodology: SA, BM, HGK, VW, ML; Investigation: SA, IA, SS, FMA, CP, BL, LLR, YK; Data analysis: SA, KB, SSA; Resources: OTK; Writing - Original Draft: SA, OTF; Writing - Review & Editing: SA, BM, HGK, ML, OTF; Supervision: BM, VW, OTK, HGK, ML, OTF; Project administration: OTF; Funding acquisition: BM, OTK, VW, HGK, ML, OTF

## Materials and Methods

### CRISPR/Cas9-based gene-editing in primary resting CD4 T cells

Gene-editing in activation-neutral CD4 T cells was performed as described previously (Albanese et al. 2022, Morath et al. 2024). Briefly, CD4 T cells were isolated by negative selection (RosetteSep Human CD4 T cells enrichment cocktail #15062, StemCell Technologies) from buffycoats obtained from healthy donors (DRK Blutspende Mannheim; ethics committee of the Medical Faculty, Heidelberg University S-024/2022). 1×10^7^ cells were washed with PBS and resuspended in 100 μl of P3 primary nucleofection buffer (P3 Primary Cell 4D-Nucleofector® X Kit L, Lonza, V4XP-3024). The cell solution was then added to a master mix of the different sgRNA-Cas9 RNP complexes (sgRNA designed and ordered from Synthego; SpCas9 nuclease from IDT #1081059) pre-prepared as described before (Albanese et al. 2022) and transferred to the large cuvettes provided in the kit. Nucleofection was performed with the program EH-100 in the Lonza 4D-Nucleofector X Unit. Cells were transferred to pre-warmed ΔRPMI (RPMI 1640, Gibco) for 10 min at 37°C and then cultured in 96-well V bottom plates (2.5×10^5^ cells/well) with supplemented RPMI (10% fetal bovine serum, 100 unites/mL penicillin, 100 mg/mL streptomycin, 2 ng/ml interleukin (IL)-7 (Peprotech, #200-7) and IL-15 (Peprotech, #200-15)). Medium change was performed every 3-4 days and successful gene-editing was validated by protein depletion after 2.5 weeks using western blot.

### Activation and HIV-1 infection of CD4 T cells

Activated CD4 T cells were generated by adding 25 μL of anti-CD3/CD28 mAb coated Dynabeads (11132D, ThermoFisher Scientific) to 1×10^6^ resting CD4 T cells for 3 days. The beads were removed by magnetic separation prior to further analyses. Knockout activated CD4 T cells were generated similarly after confirmation of protein depletion in resting CD4 T cells. HIV-1NL4-3 NLENG GFP reporter virus was produced by co-transfecting 30 μg NLENG GFP proviral plasmid (Gag p6 region modified to create a Vpx-interaction motif (Achuthan et al. 2018), with either 15 μg pcDNA 3.1 [HIV-1 - Vpx] or pcDNA 3.1 3xFlag SIVmac239 Vpx [HIV-1 +Vpx]. For virus incorporating tagged Vpr.IN.GFP, 30 μg pNL4.3 plasmid was cotransfected with 7.5 μg Vpr.IN.eGFP along with 15 μg pcDNA or Vpx plasmid in 293T cells for virus production. Infectious units of the concentrated virus were determined by performing a standard HIV-1 infectivity assay using the TZM-bl reporter cells as described previously (Ananth et al. 2024) and the number of viral particles were determined by an RT assay (SG-PERT (Pizzato et al. 2009)). For HIV-1 infection, 2×10^5^ activated or resting CD4 T cells were seeded in a 96-well plate and infected with 1×10^5^ infectious units (0.5 infectious units per cell determined on TZM-bl reporter cells) of the virus by spinoculation for 2.5 h at 650 g. Of note, the titers of virus carrying Vpx were approximately 3 to 5-fold higher than without Vpx quantified by SG-PERT. However, virus input was always normalized for infectious activity rather than particle number for the NLENG virus. For experiments with the non-infectious NNHIV (Muller et al. 2021) VLPs, 2×10^11 pU RT/ μl per 2×10^5^ cells was used. Medium exchange (with fresh IL-7/15) was performed after 24 h and the cells were harvested at 72 h to evaluate GFP expression by flow cytometry (FACS Celesta, BD Biosciences).

### Western blot

Knockout was validated in resting CD4 T cells by lysing 2×10^5^ cells in 40 μl 6X sample buffer. The lysates were separated on SDS-polyacrylamide gel electrophoresis (8-12% gels) and blotted onto nitrocellulose membranes for 1 h using the Power Blotter XL System (Invitrogen). After blocking the membranes in 5% bovine serum albumin (BSA)/ phosphate buffered saline-Tween20 (PBS-T) they were incubated with primary antibodies overnight at 4°C. After three washes in 1X PBS-T, the membranes were incubated in horseradish peroxidase (HRP)-conjugated secondary antibodies (Life Technologies) for 1 h at room temperature and visualized by chemiluminescence. The following primary antibodies were used: rabbit anti-CA (in-house, 1:1000), rabbit anti-nucleoporin p62 (ab96134, Abcam, 1:1000), rabbit anti-NUP153 [QE5] (ab24700, Abcam, 1:1000), rabbit anti-glyceraldehyde-3-phosphate dehydrogenase (GAPDH) (14C10; Cell Signaling, 1:2000), mouse anti-vinculin hVIN-1 (V9264, Sigma Aldrich, 1:2,000), rabbit recombinant anti-NUP153 (ab171074, Abcam, 1:1000), rabbit anti-RanBP2 (NUP358) (ab64276, Abcam, 1:500), rabbit anti-NUP54 (HPA035929, Atlas Antibodies, 1:1000), rabbit anti-NUP58 (HPA039360, Atlas Antibodies, 1:1000), rat anti-NUP98 [2H10] (ab50610, Abcam, 1:2000), rabbit anti-TPR (ab70610, Abcam, 1:1000), mouse anti-NUP35 (ab169162, Abcam, 1:1000).

### ddPCR and qPCR

Briefly, 2×10^5^ resting and activated CD4 T cells per well were infected with DNAse treated HIV-1 NLENG GFP virus (+/-Vpx) and two wells per condition were harvested at 48 hpi for analysis of 2-LTR circles and reverse transcribed products by ddPCR and at 72 hpi for HIV-1 transcripts by qPCR. Efavirenz and Raltegravir-treated conditions were used as control. For ddPCR, the cell pellets were lysed in 50 μl lysis buffer and 1:5 dilution was used as input. Host gene ribonuclease P protein subunit p30 (rpp30) was used as an internal loading control. All primer an probe sequences are listed below. For HIV-1 RNA quantification, total RNA was isolation using RNeasy Micro Kit, (Qiagen, #74004) kit according to manufacturer’s protocol. 250 ng μl RNA was used as input for cDNA synthesis using High-capacity cDNA Reverse Transcription Kit with RNase inhibitor (Applied Biosystems, #4374967) followed by qPCR. Primers used for qPCR were described previously by Houszet et al., (Houzet et al. 2007).

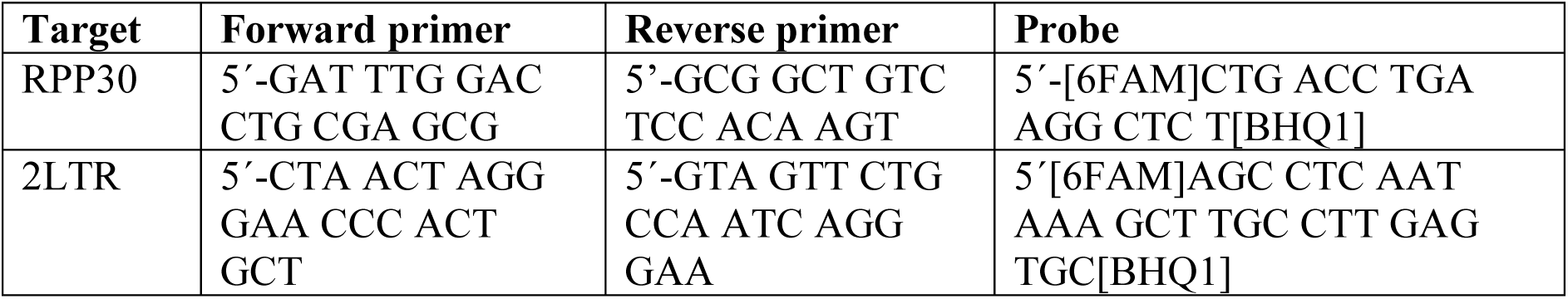
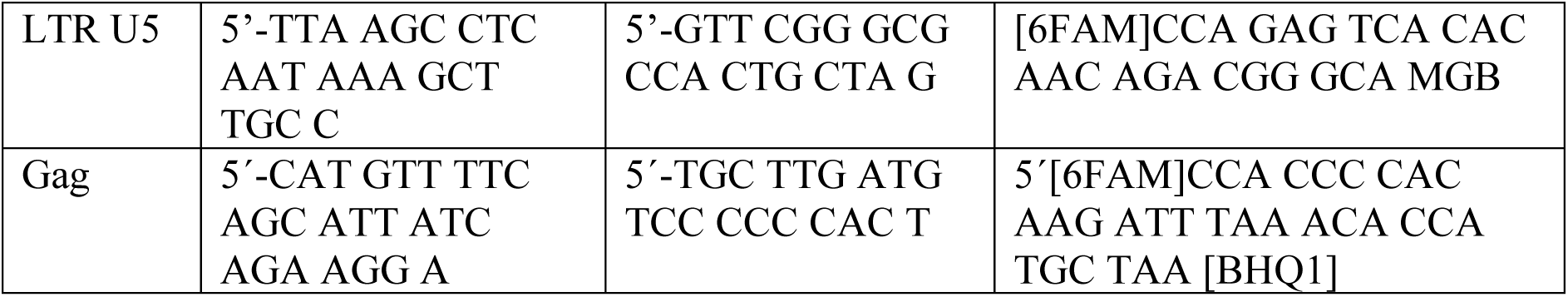

### Immunofluorescence

For microscopy-based analysis, 2×10^5^ infected resting or activated CD4 T cells were seeded onto polyethylenimine (PEI)-coated 15-well μ-Slide Angiogenesis (ibidi) for 30 minutes and subsequently fixed with 3% paraformaldehyde (PFA) for 15 min. For visualizing intracellular capsids, infected cells were resuspended in 40 μl medium containing 15 μM PF74, transferred to the imaging dish and incubated at 37°C for 1 hour. Supernatant was carefully removed and fixed with 3% PFA for 15 minutes. After fixation and subsequent washing for three times with 1X PBS, the cells were permeabilized with 0.5% Triton X-100 for 15 minutes. Cells were washed and incubated with 100% methanol for 10 minutes (for intranuclear capsid visualization) or directly blocked with 1% BSA/PBS for 30 minutes. Primary antibodies were diluted in blocking buffer, and the cells were treated for 1.5 h at room temperature. Primary antibodies used: rabbit anti-CPSF6 (#HPA039973, Atlas Antibodies, 1:200), mouse anti-SC35 (#ab11826, Abcam, 1:200), rabbit anti-CA (in house, 1:1000), mouse anti-LAMINB1 C-5 (#sc-365962, Santa Cruz Biotechnology, 1:100). After washing three times with 1X PBS, Alexa-Fluor conjugated secondary antibodies (life Technologies) were added at a dilution of 1:1000 and incubated for 45 minutes at room temperature. For super-resolution microscopy, STAR RED (STRED-1001-500UG) and STAR ORANGE (STORANGE-1002-500UG) (Abberior) secondary antibodies were used at a final dilution of 1:500. Under some conditions, the nucleus was defined using Hoechst (1:1000) by treating the samples at the final step for 2 minutes.

### Confocal and super-resolution microscopy

Multichannel Z-stack confocal images of nuclear volumes were acquired at the inverted PerkinElmer Ultra VIEW VoX 3D spinning disk confocal microscope (Perkin Elmer) using a 100X objective with a Z spacing of 200 nm. Nuclear volume was defined either by Lamin B1 or Hoechst staining. Images were acquired in the 405-, 488-, 568- and 640 nm channels depending on the fluorophore.

Airyscan point laser scanning confocal microscopy was conducted using a Zeiss LSM900 microscope (Carl Zeiss Microscopy). Z-stacks (spacing of 150 nm) across the entire nuclear volume was acquired by defining the nuclear volume as mentioned above using a 63X objective under the Airyscan super resolution mode. Zen Blue 3.1 software was used for automatic 3D processing of acquired images with default Airyscan Filtering strength.

To visualize single NPCs, Stimulated Emission Depletion (STED) super-resolution microscopy was performed using a STED system and Imspector software (Abberior Instruments GmBH). A 100X oil immersion objective was used, and images were acquired using the 590 and 640 laser lines at a pixel size of 15 nm. Depletion STED laser power was set to 10% for the 590 nm laser line and 15% for the 640 nm laser line. Half nuclear volume Z-stack images were acquired from the bottom to the center of the nucleus (defined by Lamin B1) with a Z-spacing of 150 nm.

### Nuclear permeability assay using fluorescent dextran

Fluorescent dextran of predefined molecular weight was used to determine the nuclear permeability in gene edited CD4 T cells as previously described (Raices and D’Angelo 2022). Gene-edited resting and activated CD4 T cells were seeded at a concentration of 2×10^5^ cells per 96-well in 100 μl supplemented RPMI medium containing either 20 μg/ml 3000 MW dextran-Texas Red (#D3328, Invitrogen) or 20 μg/ml of 70000 MW dextran-Texas Red (#D1830, Invitrogen). The cells were incubated at 37 °C for 1 hour to allow for dextran uptake. After the incubation, the medium was aspirated, and the cells were resuspended in 50 μl 1X PBS and seeded onto poly-L-lysine coated 10 mm coverslips. After 10 minutes of settling time, the cells were fixed with 3% PFA for 15 minutes and 0.01% Triton-×100 for 5 minutes. After washing three times with 1X PBS, Hoechst staining was performed to define the nucleus. The dye was washed, and the coverslips were mounted using Mowiol. Z-stack images were obtained using a confocal spinning-disk microscope (Perkin Elmer).

### Image analyses and visualization

Images were analyzed using Fiji (Schindelin et al. 2012) and intranucleoplasmic HIV-1 complexes were quantified as described in detail before (Ananth et al. 2024). Briefly, HIV-1 CA foci within the nucleus were manually analyzed across the entire nuclear volume by drawing line profiles to evaluate their intensities in the CA and LAMINB1 channels from raw images to determine their localization. Capsid signals very close to the nuclear boundary or at the nuclear envelope, indicated by an overlapping intensity with LAMINB1 were eliminated from the analyses. Manual scoring was performed per cell, and the % nuclear import was determined by taking the % of cells that contain at least one intranucleoplasmic capsid foci and the total cells analyzed in the sample.

Nuclear localization of fluorescent dextran was quantified by drawing line profiles of 3 μm from the inside of the nucleus to the extracellular space. Signal intensities of dextran over 3000 A.U (arbitrary units) within the nucleus that overlapped with Hoechst intensities were scored as cells that show nuclear permeability to dextran.

Images from super-resolution microscopy were analyzed through a pipeline that involved deconvolution using Huygens Professional, segmentation using Ilastik and 3D statistics using Imaris. Briefly, images from the STED were imported into Huygens Professional software and the microscopy parameters (numerical aperture: 1.4, backprojected pinhole: 250, excitation wavelength: 561 nm and 640 nm, emission wavelength: 615 nm and 666 nm, excitation fill factor: 1.38, STED depletion mode: pulsed, STED wavelength: 775 nm, STED saturation factor: 20, STED immunity fraction: 5%) were entered. The preset deconvolution wizard was used with automatic background estimation and stabilization. Images were exported as 16-bit .TIFF files per channel that were then loaded into Ilastik. All images per channel per donor were loaded at the same time into Ilastik for segmentation and 4-5 images were chosen for training. Separate training and segmentation were performed for the different donors to correct for staining discrepancies, especially in the LAMINB1 channel between donors. Segmented images were opened in Imaris and 3D cropped in Z-axis to ensure only a thin layer of LAMINB1 (approx. 3-4 Z-slices) is chosen for further analyses. Surface rendering was performed using the images from LAMINB1 channel and ‘Spots’ option was used to detect punctate NPC signals. Spots in the NUP channel with size < 0.192 were filtered out. NPCs at the nuclear envelope were analyzed by filtering signals using the ‘shortest distance to surface’ option relative to the LAMINB1 surface. NPC spots within 0.0339 μm from the LAMINB1 surface were considered to be located at the nuclear envelope and ‘distance to nearest neighbor’ analyses were performed.

### Soft X-ray tomography

Full-rotation SXT was performed following the protocol described by Chen et al (Chen et al. 2022). Tomographic reconstruction was carried out using AREC3D. For post-processing, an unsharp mask was applied to enhance contrast, using a radius (sigma) of 15 pixels and a mask weight of 0.60. Levels of X-ray attenuation within the nucleus was represented as linear absorption coefficient [LAC] values.

### Integration sites (ISs) sequencing

Integration sites were sequenced from HIV-1 infected NT control or Nup62 KO resting and activated CD4 T cells harvested at 3 days post-infection (dpi). Genomic DNA was extracted using the Qiagen Blood & Tissue Kit following the manufacturer’s instructions. Ligation-mediated PCR (LM-PCR) and next-generation sequencing were performed as previously described with modification (Serrao et al. 2016). Briefly, genomic DNA (1.6–2.64 µg, kept constant between samples from the same donor) was sheared to approximately to 500 bp using a Covaris S220 Ultrasonicator. Fragmented DNA was end-repaired (Thermo Fisher Fast DNA End Repair Kit), A-tailed (NEBNext® dA-Tailing Module), and purified after each step using the Macherey-Nagel PCR Purification Kit. An asymmetric double-stranded linker was ligated overnight at 12 °C using 1.5 µM linker and 800 U T4 ligase (NEB) and purified with 0.9× AMPure XP beads (Beckman Coulter). The first PCR was performed in parallel reactions using 20-30 ng of ligation products with 0.8 µM 3’ LTR- and linker-specific primers under the following conditions: 94 °C 2 min; 30 cycles of 94 °C 15 s, 64 °C 30 s, 72 °C 30 s with final extension at 72 °C 10 min. Products were purified using 0.9× AMPure XP beads. A nested PCR using 0.8 µM inner HIV-1 LTR and the same linker primer containing Illumina adapters and unique dual indexes (UDI) was performed for 6 cycles under identical conditions, followed by purification with 0.9× AMPure XP beads. Library quality was assessed using an Agilent Bioanalyzer and quantified with the NEBNext® Library Quant Kit for Illumina®. Sequencing was performed at the c.ATG Core Facility (University of Tübingen) on an Illumina MiSeq platform (2×150 bp). The list of HIV-1 integrations is available at: https://zenodo.org/records/24394467

### Bulk RNA sequencing

For RNAseq analysis, at least 1×10^6^ infected activated and resting NT and NUP62 KO cells were harvested and processed for RNA isolation as mentioned before for PCR analysis. 200 ng RNA was prepared per sample per condition and sent for library construction and sequencing using Illumina NovaSeq X high throughput sequencing. The reads obtained were mapped using HISAT2 software to the reference human genome (Homo_sapiens.GRCh38_release95.genome.fa.). Mapped reads were assembled using StringTie contrast pairs for subsequent differential gene expression (DEG) analysis. DEGs between conditions were analyzed by DESeq2 using the following thresholding criteria: fold changes (FC) ≥ 1.5 and a p-value < 0.05 and plotted using Prism 10 (GraphPad). The raw sequencing files can be obtained from: https://zenodo.org/records/24354631.

### Bioinformatic analysis

#### SPIN state analysis from IS sequencing

Raw paired-end sequencing reads were processed using the integration site mapping pipeline used before (Penzo et al. 2025). Adapter and viral long terminal repeat (LTR) sequences were trimmed and reads containing verified LTR–host junctions were aligned to the human reference genome (hg38). Uniquely mapped reads were filtered for alignment quality. To account for genomic mappability and fragmentation biases, matched in silico random control datasets were generated across the hg38 genome using the random genome fragment simulation framework described by Greg Bedwell (Random_insertions.R). Experimental integration coordinates from CD4 T cells and simulated random controls were then intersected with Spatial Position Inference of the Nuclear (SPIN) genome state annotations (Speckle, Interior_Act1, Interior_Act2, Interior_Act3, Interior_Repr1, Interior_Repr2, Near_Lamina1, Near_Lamina2, and Lamina) generated for K562 cells (Gholamalamdari et al. 2025). Integration sites in bins without a SPIN assignment, and sites outside the annotated regions, were excluded (0.5–1.4% and 3.1–3.7% of sites, respectively); proportions were calculated across the nine remaining states. Integration targeting preferences across each SPIN compartment were quantified by calculating the proportion and \(\log _{2}\) fold-enrichment of unique HIV-1 integration sites relative to matched random controls.

#### RNAseq read processing and alignment

Paired-end RNA-seq reads were assessed for quality with FastQC (version 0.12.1) (Andrews 2010). Adapter removal and quality trimming were performed with Trim Galore (version 0.6.1) (F Krueger), using a Phred quality cutoff of 5 and a minimum adapter overlap of 5 bp. Trimmed reads were aligned to the human genome (GRCh38/hg38) with STAR (version 2.7.11b) (Dobin et al. 2013) in unstranded splice-aware mode against the GENCODE annotation (release 50) (Mudge et al. 2025), permitting intronic gaps between 20 bp and 1 Mb.

#### Gene-level counts and TPM

Uniquely mapped reads were assigned to genes with featureCounts (Subread, version 2.1.1) (Liao et al. 2014) at the exon level and summarized per gene in unstranded mode using the same GENCODE annotation. For all expression analyses, the gene universe was restricted to protein-coding genes.

#### Expression-integration analyses

Because the RNA-seq and integration-site donors were distinct individuals, each gene was assigned the mean expression across all RNA-seq libraries of a given activation state. The relationship between gene expression and integration was summarized by ranking expressed genes (TPM > 1) into deciles of expression within each activation state. Per each decile, the fraction of genes targeted by at least one integration and the density of integration sites per Mb of gene body were computed. Because non-targeting and edited samples differed in the total number of recovered sites, the analysis was repeated after rarefaction. Briefly, within each activation state, the genotype with more sites was randomly downsampled to match the smaller, the decile metrics were recomputed, and the results were summarized over 200 draws.

#### Integration into silent genes

To assess targeting of genes below the expression threshold, each genic integration site was classified by the expression of its host gene in the pooled reference as silent (TPM ≤ 1) or expressed (TPM > 1). For every sample the fraction of integration sites in silent genes was computed, and this fraction was compared between KO62 and NT within each activation state using the paired t-test.

#### Quantification of viral transcripts

To quantify expression of the integrated provirus, RNA-seq reads were aligned with STAR to a combined reference comprising the human genome (GRCh38/hg38) and the HIV proviral construct. Viral-feature counts were normalized together with the host-gene counts within DESeq2. Normalized viral-gene expression was compared between KO62 and NT within each activation state, with samples paired by donor.

#### Locus-level knockout verification

NUP62 knockout was examined at nucleotide resolution across the NUP62 locus (final exon, chr19:49,908,500-49,911,500). Per-base RNA-seq coverage over the locus was computed and normalized to counts per million (CPM) using bamCoverage (DeepTools, version 3.5.6)(Ramirez et al. 2016). All expression analyses were performed using R (version 4.5.2) https://www.R-project.org/.

### Statistical analysis

Statistical analysis was performed using Prism 10 (GraphPad). For donor-matched comparisons between NT and KO samples, paired ratio t test was used. Parametric or non-parametric analysis was performed based on the test for normality. One sample t test was used to calculate significance where control samples were normalized to 1. All statistical tests used are mentioned in the respective figure legends.

SPIN state distributions were compared between NUP62 KO and NT cells by χ² tests of independence across the nine states. Per-state comparisons used Fisher’s exact test with Holm correction for multiple testing and are reported as odds ratios with 95% confidence intervals. Whether the NUP62 KO effect depended on activation state was tested as the ratio of odds ratios for the interior-active states (Speckle and Interior_Act1–3) between resting and activated cells. These analyses were performed in Python (SciPy).

