## Supplementary file for "NUP62 acts as T cell activation-dependent switch to regulate post nuclear import steps of HIV-1 replication in primary CD4 T cells"

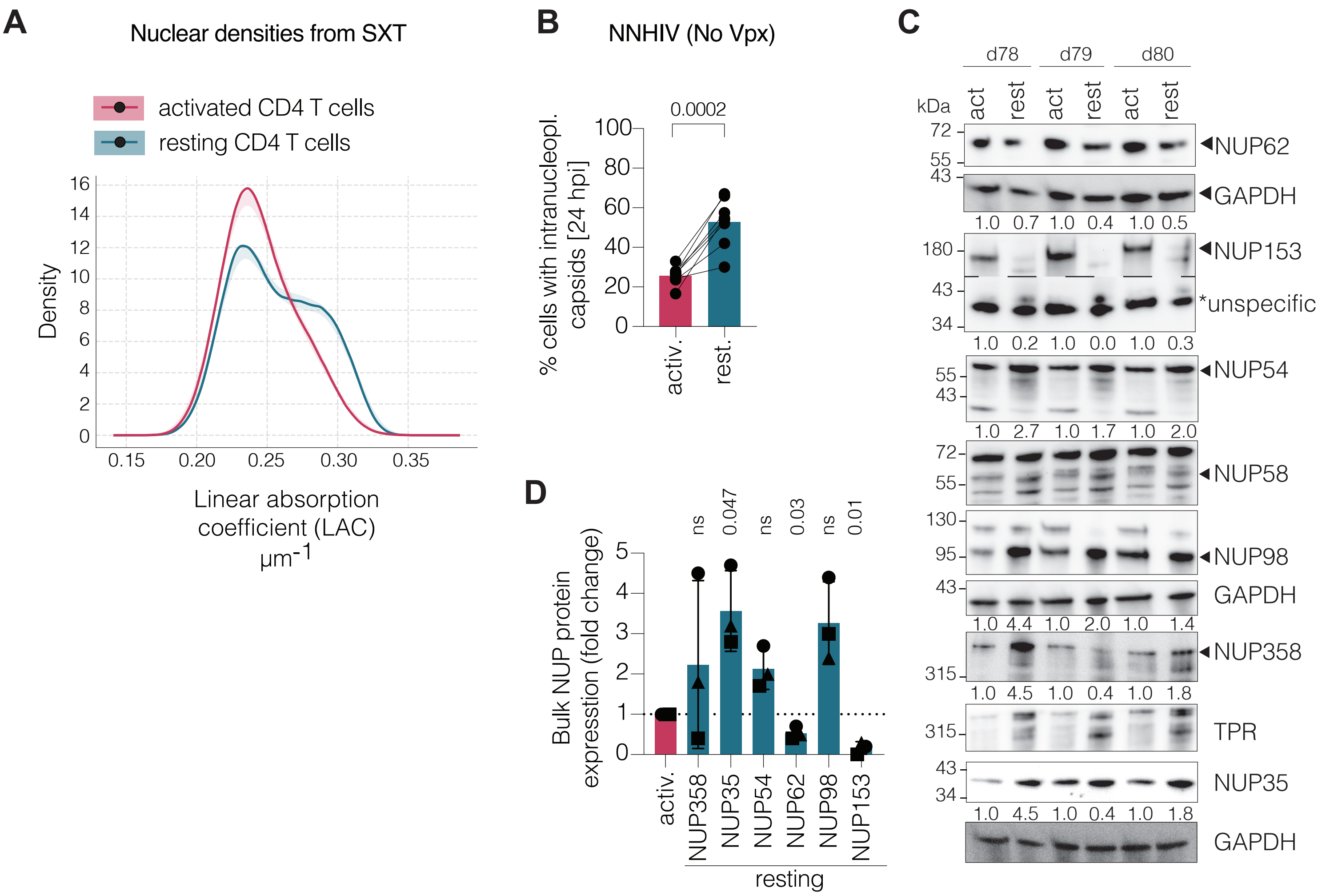
A
Nuclear densities from SXT
activated CD4 T cells
resting CD4 T cells
Density
0
2
4
6
8
10
12
14
16
0.15
0.20
0.25
0.30
0.35
Linear absorption coefficient (LAC) µm-1
B
NNHIV (No Vpx)
0.0002
% cells with intranucleopl. capsids [24 hpi]
0
20
40
60
80
100
activ.
rest.
C
d78
d79
d80
kDa
act
rest
act
rest
act
rest
72
55
43
NUP62
GAPDH
1.0 0.7 1.0 0.4 1.0 0.5
180
43
34
NUP153
*unspecific
1.0 0.2 1.0 0.0 1.0 0.3
55
43
NUP54
1.0 2.7 1.0 1.7 1.0 2.0
72
55
NUP58
130
95
NUP98
43
GAPDH
1.0 4.4 1.0 2.0 1.0 1.4
315
NUP358
1.0 4.5 1.0 0.4 1.0 1.8
315
TPR
43
34
NUP35
1.0 4.5 1.0 0.4 1.0 1.8
GAPDH
D
Bulk NUP protein expression (fold change)
0
1
2
3
4
5
ns
0.047
ns
0.03
ns
0.01
activ.
NUP358
NUP35
NUP54
NUP62
NUP98
NUP153
resting

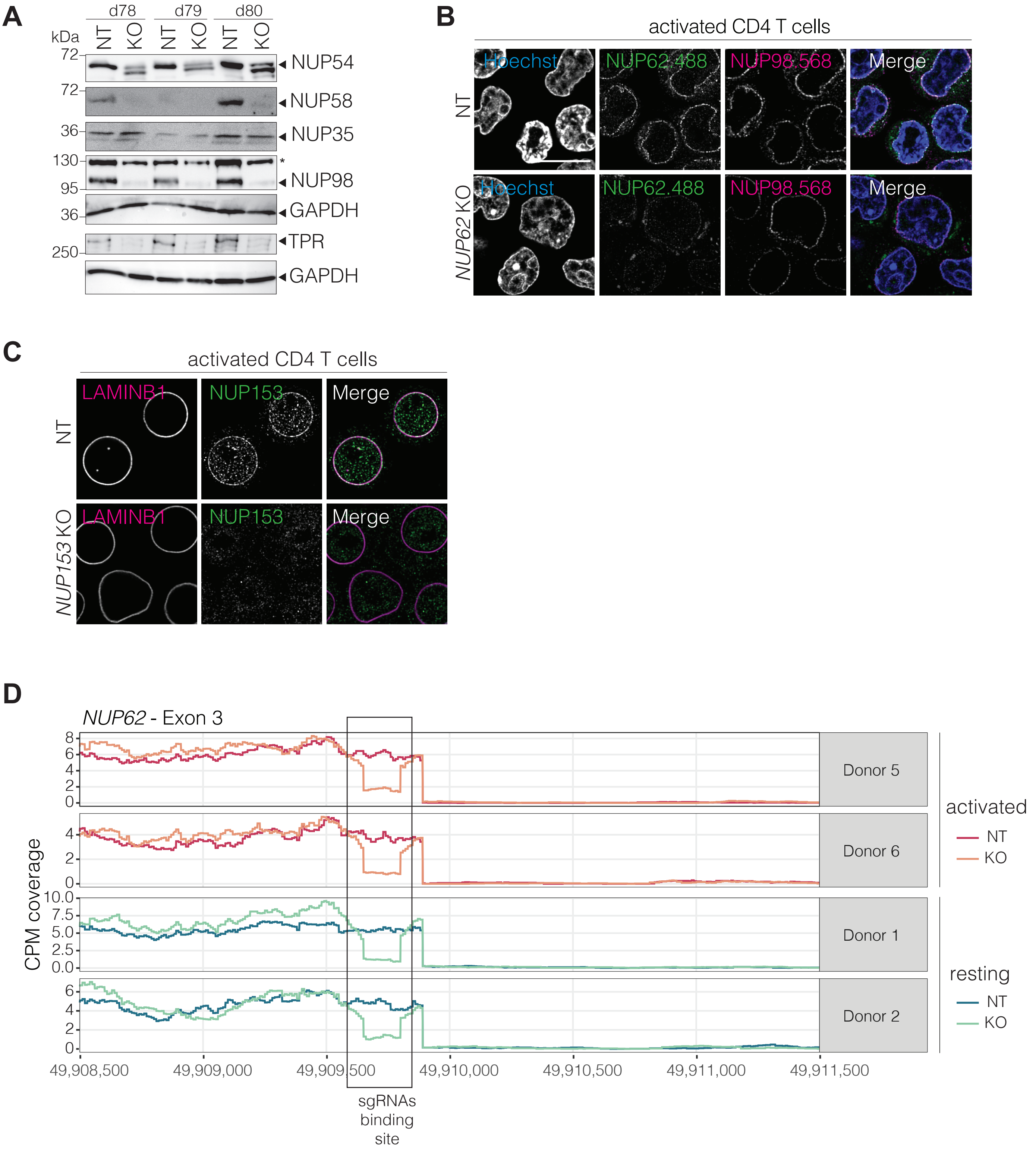
A
d78
d79
d80
kDa
NT
KO
72
NUP54
72
NUP58
36
NUP35
130
*
NUP98
95
36
GAPDH
TPR
250
GAPDH
B
activated CD4 T cells
Hoechst
NUP62.488
NUP98.568
Merge
NT
NUP62 KO
C
activated CD4 T cells
LAMINB1
NUP153
Merge
NT
NUP153 KO
D
NUP62 - Exon 3
CPM coverage
Donor 5
Donor 6
Donor 1
Donor 2
activated
NT
KO
resting
NT
KO
49,908,500
49,909,000
49,909,500
49,910,000
49,910,500
49,911,000
49,911,500
sgRNAs binding site

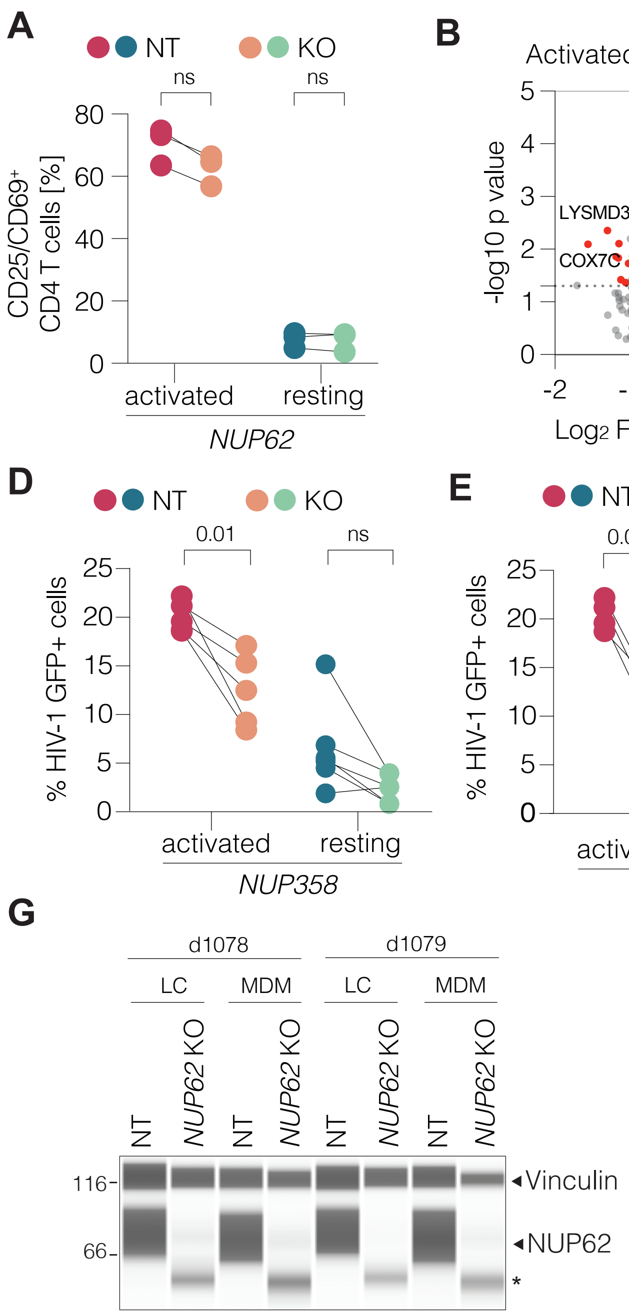
A
NT
KO
ns
ns
CD25/CD69+ CD4 T cells [%]
activated
resting
NUP62
B
Activated (NUP62 KO vs NT)
-log10 p value
LYSMD3
COX7C
PCDHB7
Log2 FC (act6h KO vs NT)
C
Resting (NUP62 KO vs NT)
-log10 p value
STRBP
GLCCI1
Log2 FC (resting KO vs NT)
D
NT
KO
0.01
ns
% HIV-1 GFP+ cells
activated
resting
NUP358
E
NT
KO
0.006
ns
% HIV-1 GFP+ cells
activated
resting
NUP153
F
NT
KO
+ Vpx
- Vpx
% HIV-1 GFP+ cells
activated
resting
NUP62
G
d1078
d1079
LC
MDM
LC
MDM
NT
NUP62 KO
NT
NUP62 KO
NT
NUP62 KO
NT
NUP62 KO
116
66
Vinculin
NUP62
*

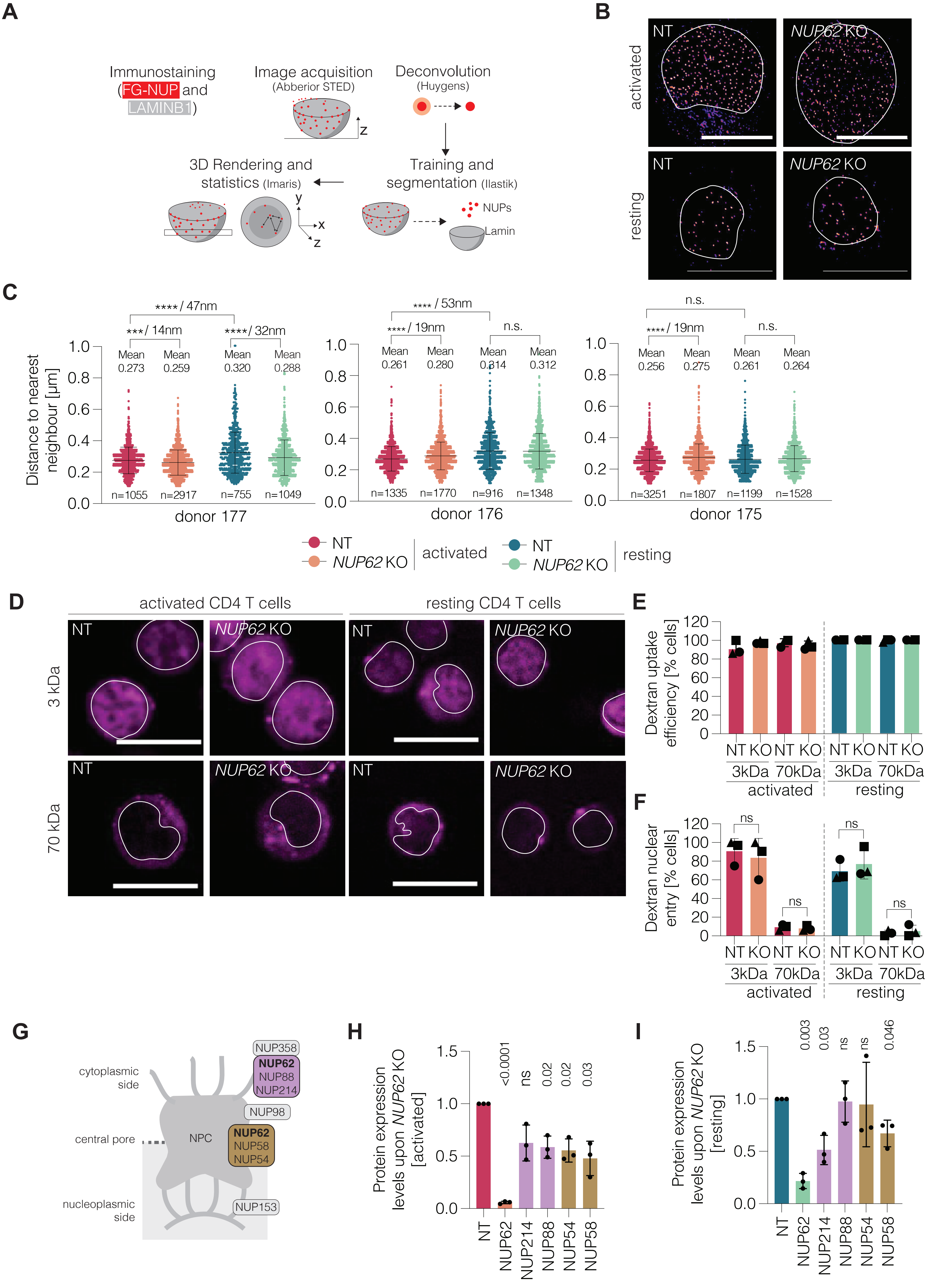
A
Immunostaining (FG-NUP and LAMINB1)
Image acquisition (Abberior STED)
z
Deconvolution (Huygens)
Training and segmentation (Ilastik)
NUPs
Lamin
3D Rendering and statistics (Imaris)
y
x
z
B
NT
NUP62 KO
activated
resting
C
Distance to nearest neighbour [µm]
****/ 47nm
***/ 14nm
****/ 32nm
Mean 0.273
Mean 0.259
Mean 0.320
Mean 0.288
n=1055
n=2917
n=755
n=1049
donor 177
**** / 53nm
****/ 19nm
n.s.
Mean 0.261
Mean 0.280
Mean 0.314
Mean 0.312
n=1335
n=1770
n=916
n=1348
donor 176
n.s.
****/ 19nm
n.s.
Mean 0.256
Mean 0.275
Mean 0.261
Mean 0.264
n=3251
n=1807
n=1199
n=1528
donor 175
NT
NUP62 KO
activated
NT
NUP62 KO
resting
D
activated CD4 T cells
resting CD4 T cells
3 kDa
70 kDa
NT
NUP62 KO
E
Dextran uptake efficiency [% cells]
NT KO NT KO NT KO NT KO
3kDa 70kDa 3kDa 70kDa
activated
resting
F
Dextran nuclear entry [% cells]
ns
ns
ns
ns
NT KO NT KO NT KO NT KO
3kDa 70kDa 3kDa 70kDa
activated
resting
G
cytoplasmic side
central pore
nucleoplasmic side
NPC
NUP358
NUP62
NUP88
NUP214
NUP98
NUP62
NUP58
NUP54
NUP153
H
Protein expression levels upon NUP62 KO [activated]
<0.0001
ns
0.02
0.02
0.03
NT
NUP62
NUP214
NUP88
NUP54
NUP58
I
Protein expression levels upon NUP62 KO [resting]
0.003
0.03
ns
ns
0.046
NT
NUP62
NUP214
NUP88
NUP54
NUP58

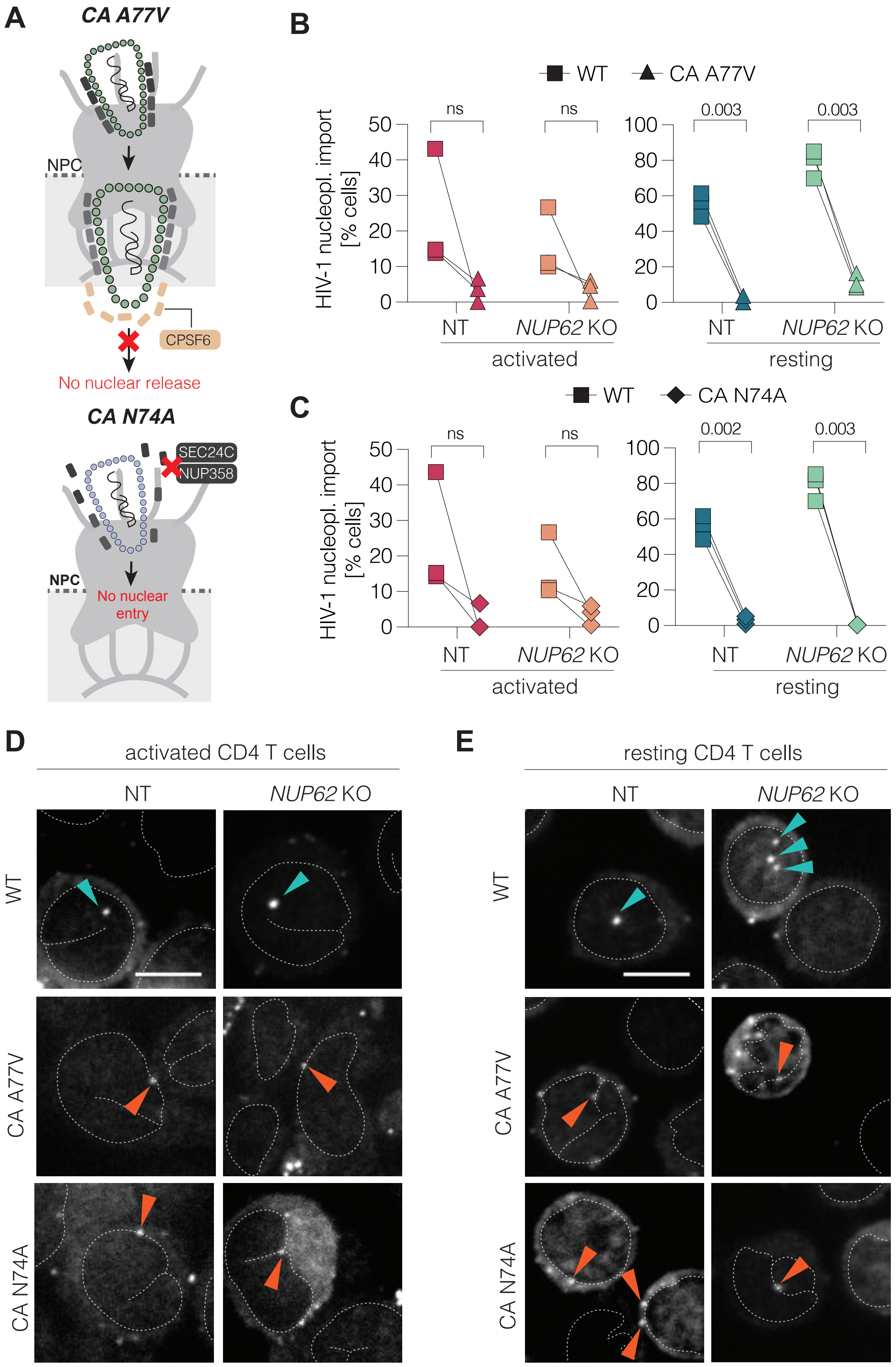
A
CA A77V
NPC
CPSF6
No nuclear release
CA N74A
SEC24C
NUP358
NPC
No nuclear entry
B
WT
CA A77V
HIV-1 nucleopl. import [% cells]
ns
ns
0.003
0.003
NT
NUP62 KO
activated
NT
NUP62 KO
resting
C
WT
CA N74A
HIV-1 nucleopl. import [% cells]
ns
ns
0.002
0.003
NT
NUP62 KO
activated
NT
NUP62 KO
resting
D
activated CD4 T cells
NT
NUP62 KO
WT
CA A77V
CA N74A
E
resting CD4 T cells
NT
NUP62 KO
WT
CA A77V
CA N74A

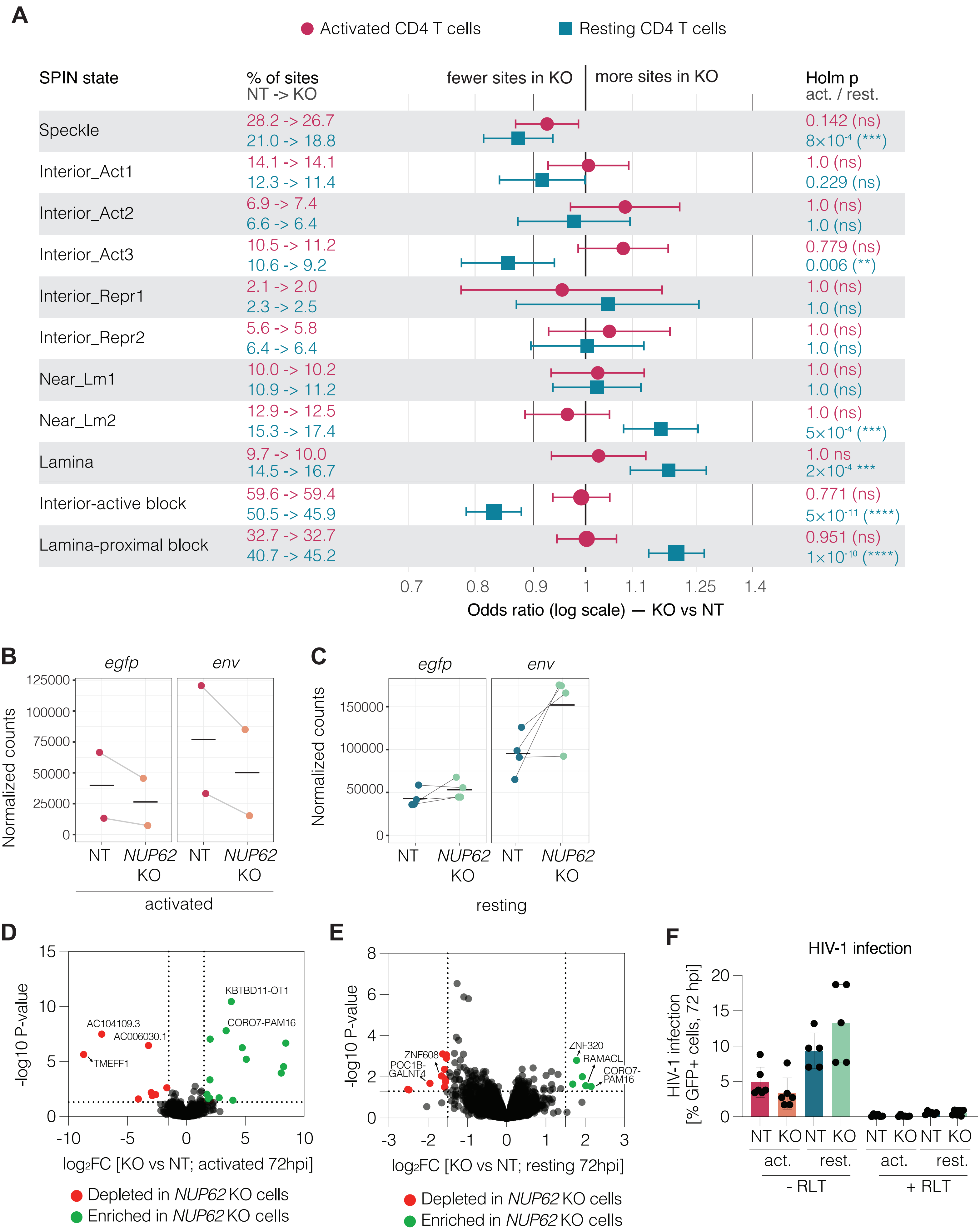
A
Activated CD4 T cells
Resting CD4 T cells
SPIN state
% of sites NT -> KO
fewer sites in KO
more sites in KO
Holm p act. / rest.
Speckle 28.2 -> 26.7 21.0 -> 18.8 0.142 (ns) 8×10-4 (***)
Interior_Act1 14.1 -> 14.1 12.3 -> 11.4 1.0 (ns) 0.229 (ns)
Interior_Act2 6.9 -> 7.4 6.6 -> 6.4 1.0 (ns) 1.0 (ns)
Interior_Act3 10.5 -> 11.2 10.6 -> 9.2 0.779 (ns) 0.006 (**)
Interior_Repr1 2.1 -> 2.0 2.3 -> 2.5 1.0 (ns) 1.0 (ns)
Interior_Repr2 5.6 -> 5.8 6.4 -> 6.4 1.0 (ns) 1.0 (ns)
Near_Lm1 10.0 -> 10.2 10.9 -> 11.2 1.0 (ns) 1.0 (ns)
Near_Lm2 12.9 -> 12.5 15.3 -> 17.4 1.0 (ns) 5×10-4 (***)
Lamina 9.7 -> 10.0 14.5 -> 16.7 1.0 ns 2×10-4 ***
Interior-active block 59.6 -> 59.4 50.5 -> 45.9 0.771 (ns) 5×10-11 (****)
Lamina-proximal block 32.7 -> 32.7 40.7 -> 45.2 0.951 (ns) 1×10-10 (****)
0.7 0.8 0.9 1 1.1 1.25 1.4
Odds ratio (log scale) — KO vs NT
B
egfp
env
Normalized counts
125000 100000 75000 50000 25000 0
NT NUP62 KO NT NUP62 KO
activated
C
egfp
env
Normalized counts
150000 100000 50000 0
NT NUP62 KO NT NUP62 KO
resting
D
-log10 P-value
15 10 5 0
KBTBD11-OT1
CORO7-PAM16
AC104109.3
AC006030.1
TMEFF1
-10 -5 0 5 10
log2FC [KO vs NT; activated 72hpi]
Depleted in NUP62 KO cells
Enriched in NUP62 KO cells
E
-log10 P-value
8 6 4 2 0
ZNF320
RAMACL
CORO7-PAM16
ZNF608
POC1B-GALNT4
-3 -2 -1 0 1 2 3
log2FC [KO vs NT; resting 72hpi]
Depleted in NUP62 KO cells
Enriched in NUP62 KO cells
F
HIV-1 infection
HIV-1 infection [% GFP+ cells, 72 hpi]
20 15 10 5 0
NT KO NT KO NT KO NT KO
act. rest. act. rest.
- RLT
+ RLT

**Supplementary Figure 1. Differences between resting and activated CD4 T cells. (A)** Nuclear LAC values depicted as a probability density, indicating how frequently different linear attenuation coefficient LAC values occur within the nucleus in the y-axis and the exact LAC values in the x-axis. **(B)** Nuclear import efficiency of NNHIV (integration-deficient HIV-1 $NL_{4-3}$). Resting and autologous anti-CD3/CD28-activated CD4 T cells were spin-infected with $2 \times 10^{10}$ pU RT/ml of NNHIV. At 24 hpi, cells were harvested, immunostained for the viral capsid protein, and quantified as described in panel 1F. Paired t test between activated and resting CD4 T cells were performed to calculate statistical significance. **(C)** Western blots showing the expression of various NUPs and TPR in activated and resting CD4 T cells from three independent donors. GAPDH was used as a loading control. Values below the blots represent densitometric quantifications, normalized first to their respective GAPDH bands and then to the expression levels in activated CD4 T cells, which were arbitrarily set to 1. **(D)** Densitometric quantifications from panel A plotted as a bar graph. Statistical significance was calculate using one sample t-test. ns: not significant.

**Supplementary Figure 2. Expression of nucleoporins and validation of knockout efficiency. (A)** Validation of *NUP54*, *NUP58*, *NUP35*, *NUP98*, and *TPR* KO in resting CD4 T cells at 2–3 weeks post-nucleofection, as assessed by western blot. Representative blots from three independent donors are shown. **(B)** Representative confocal micrographs of NT and *NUP62* KO activated CD4 T cells immunostained with antibodies against NUP62 and NUP98. Hoechst was used to stain the nucleus. Scale bar: 10 µm. **(C)** Representative Airyscan images of NT and *NUP153* KO activated CD4 T cells immunostained for NUP153. In KO cells, a marked reduction in NPC-associated NUP153 signal overlapping with LAMINB1 was observed. **(D)** RNA-seq read coverage at the *NUP62* gene locus (Exon 3) in NT and *NUP62* KO activated and resting CD4 T cells. The highlighted region indicates the binding sites of the two sgRNAs designed to target the gene. Data are shown from two independent donors.

**Supplementary Figure 3. HIV-1 infection and impact of *NUP62* KO on T cell activation and cellular transcriptome.** (**A**) Verification of T cell activation status in activated and resting CD4 T cells upon *NUP62* KO. Cells were stained with fixable viability dye for 20 min at room temperature and measured by flow cytometry, Unstained cells were gated as viable population. **(B, C)** Volcano plot comparing the microarray-based transcriptional profiles of of NT and *NUP62* KO cells under 6 h-activated (B) and resting (C) conditions. Significantly upregulated genes are highlighted in green and downregulated genes in red. **(D, E)** Infection rates, measured as the percentage of GFP-expressing cells by flow cytometry at 72 hpi, in NT and KO activated and resting CD4 T cells for the targets *NUP358* (D) and *NUP153* (E). Five or more independent donors were analyzed. **(F)** Infection of NT and *NUP62* KO cells with virus carrying Vpx for activated and no Vpx for resting conditions. **(G)** Representative virtual immunoblots from Jess indicating NUP62 protein depletion. Vinculin was used as loading control. Statistical significance was calculated using ratio paired t test for panels A, D and E.

**Supplementary Figure 4. Characterization of NT and NUP62 KO cells for NPC distribution, permeability and NUP62 interaction partners. (A)** Schematic workflow for the analysis of inter-NPC distances in activated and resting CD4 T cells. NT and *NUP62* KO cells were immunostained with FG-NUP to visualize all NPCs and LAMINB1 to delineate the nuclear boundary. STED images of the basal nuclear hemisphere were acquired along the Z-axis. Exported images were deconvolved using Huygens software and segmented using Ilastik. Segmented images were then processed in Imaris for 3D reconstruction. NPC spots overlapping with the LAMINB1 signal were subjected to nearest neighbor analysis to quantify inter-NPC distances. **(B)** Representative single Z-slice STED images of NT and *NUP62* KO resting and activated CD4 T cells stained for NPCs using an anti-FG-NUP antibody. The nuclear boundary is indicated in white, defined by LAMINB1 staining. Scale bar: 10 µm. **(C)** Nearest neighbor distance analysis performed in 3D using Imaris for nuclear NPC signals in activated and resting CD4 T cells from three independent donors. Statistical significance was determined using unpaired t-test comparing NT and KO conditions. **(D–F)** Nuclear permeability assay. (D) Representative single Z-slice confocal images of gene-edited resting and activated CD4 T cells incubated with 3 kDa and 70 kDa fluorescent dextran (Texas Red; magenta). The nuclear boundary is indicated in white, defined by Hoechst staining. Scale bar: 10 µm. (E) Frequency of cells showing overall dextran uptake (fluorescence observed in the cytoplasm). (F) Frequency of cells showing nuclear dextran entry (fluorescence observed in the nucleus). **(G)** Schematic representation of NUP62 interaction partners within the NPC: NUP88 and NUP214 on the cytoplasmic face, and NUP58 and NUP54 within the inner ring. **(H, I)** Densitometric quantification of the expression levels of the NUPs shown in (G) in activated and resting NT and *NUP62* KO CD4 T cells. Statistical significance was determined using a one-sample t-test.

**Supplementary Figure 5. Viral determinants for nuclear import remain unchanged in the absence of NUP62 in CD4 T cells. (A)** Schematic representation of HIV-1 capsid entry through the nuclear pore complex (NPC). The A77V capsid mutation permits nuclear entry but prevents release into the nucleoplasm due to loss of interaction with CPSF6 whereas N74A prevents CA interaction with several host cell factors such as NUP358 and SEC24C and prevent nuclear entry. **(B, C)** HIV-1 nuclear import in activated and resting CD4 T cells infected with wild-type (WT) or CA A77V mutant (B) or CA N74A mutant (C). (D, E) Representative confocal images of activated and resting CD4 T cells for quantification shown in B, C. Capsid staining is shown in cyan, and LAMINB1 nuclear envelope staining is shown in white. White arrows indicate capsids in the nucleoplasmic space, and orange arrows indicate capsids at the nuclear envelope. Statistical significance was determined using a paired t-test to compare NT or KO conditions.

**Supplementary Figure 6. Integration-site sequencing and transcriptomic profiles of HIV-1-infected activated and resting *NUP62* KO CD4 T cells (A)** Effect of *NUP62* KO on HIV-1 integration depicted as a forest plot. Odds ratio (NT vs KO) are plotted with 95% confidence interval. Block rows are summary contrasts (each block vs all other states). Sites pooled across donors and treated as independent. Fisher's exact, Hom-corrected within each cell state. **(B, C)** Total RNA isolated from HIV-1-infected activated and resting NT and *NUP62* KO CD4 T cells was subjected to bulk RNA sequencing at 72 h post-infection (hpi). Normalized read counts for *egfp* and the viral *env* gene in infected activated CD4 T cells from two donors (B) and resting CD4 T cells from four donors (C). **(D, E)** Differential gene-expression profiles associated with *NUP62* KO in HIV-1-infected activated (D) and resting (E) $CD4^+$ T cells at 72 hpi. Differentially expressed genes are highlighted (upregulated in green, downregulated in red. **(F)** HIV-1 infection rates in NT and *NUP62* KO activated and resting CD4 T cells, with or without Raltegravir (RLT) treatment at 72 hpi that were used for quantifying viral RNA transcripts.

**Supplementary Table 1: Frequencies of HIV-1 particles after cell entry that complete different steps in HIV-1 infection**

**(i) Activated CD4 T cells**

| Steps in HIV-1 infection | % entered particles | References |
|---|---|---|
| Nuclear Import | NA | Many studies visualized CA in the nucleus in activated CD4 T cells |

| | | |
|---|---|---|
| | | but no direct quantification of nuclear import frequency (% cells with nuclear CAs) |
| Reverse Transcription | ~50% | (Zhou *et al.* 2005) |
| Integration | ~25% (20-50% of RT products) | (Agosto *et al.* 2007) |
| Viral gene expression | ~3.75% (~15% of integrated products) | (Agosto *et al.* 2018, Santos *et al.* 2023) |

**(ii) Resting CD4 T cells**

| **Steps in HIV-1 infection** | **% entered particles** | **References** |
|---|---|---|
| Nuclear Import | ~60% | (Ananth *et al.* 2024) |
| Reverse Transcription | ~6% | (Swiggard *et al.* 2005, Baldauf *et al.* 2012, Descours *et al.* 2012) |
| Integration | ~1.2% (20 of complete RT products) | (Swiggard *et al.* 2005, Agosto *et al.* 2007) |
| Viral gene expression | ~0.12% (~10% of integrated products) | (Lassen *et al.* 2004) |

**Supplementary Table 2: sgRNA sequences for CRISPR/Cas9-based gene-editing**

| **Target/ sgRNA number** | **Sequence** |
|---|---|
| NT | GCACUACCAGAGCUAACUCA |
| NUP35_sgRNA1 | GCUGGUACAACUGGUUGUGG |
| NUP35_sgRNA4 | GGAUCUGAACCAAUGAUGCU |
| NUP35_sgRNA6 | AUUCACCUACCUGCAAGUAA |
| NUP54_sgRNA1 | UUUGGUGGUACUCAGAACAA |
| NUP54_sgRNA2 | ACUGGUUUAGGUACUGGUUU |
| NUP54_sgRNA3 | GGGACAACAUCUACAACUGC |
| NUP58_sgRNA1 | UUAAAUAAGCAACCCUUCUG |
| NUP58_sgRNA2 | UCCUCCUAGAGUGAACCCAG |
| NUP58_sgRNA3 | AUAGCAACAACUAUAACUAC |
| NUP62_sgRNA1 | CAGAGUCCGGAGCCAUGAGC |
| NUP62_sgRNA2 | GCCGGUGGAAGGGGUACUUG |
| NUP98_sgRNA1 | CACUUACUGCCAAAGCCAGU |
| NUP98_sgRNA2 | UGUGCUUGCAGUUCCAAACA |
| NUP98_sgRNA3 | AGCUGGCUGGCUAAAUGAAC |
| TPR_sgRNA1 | CCCGAGAGUGUCAAAGCUUG |
| TPR_sgRNA2 | AGAAAUAUCUCUUACCUGAA |
| TPR_sgRNA3 | UUCUUGAGAUAGUCUCUCAU |
| NUP358_sgRNA1 | GAAUCUAAGACUAUAGAAUU |
| NUP358_sgRNA2 | UAUCCUGGUUCUGGCAAAGG |

| | |
|---|---|
| NUP358_sgRNA3 | GGCGAUAGAAUUCCCGUUGC |
| NUP153_sgRNA1 | UCAACAGGGCAUUCUUAGCA |
| NUP153_sgRNA2 | CUGUUCAACAGACACAAGCG |
| NUP153_sgRNA3 | GAGAGCUCUAAUAUUACUGA |

**Supplementary Table 3: SPIN states for Activated and resting NT and *NUP62* KO cells with matching random and matching covariate controls, corrected per GC content**

| | **Interior_Act1** | **Interior_Act2** | **Interior_Act3** | **Interior_Repr1** | **Interior_Repr2** |
|---|---|---|---|---|---|
| Activated_RMC_covariate | 8670 | 1560 | 2093 | 2104 | 2334 |
| Activated_RMC_rand | 3409 | 1616 | 2889 | 1221 | 2650 |
| Resting_RMC_covariate | 8612 | 1552 | 2268 | 1979 | 2310 |
| Resting_RMC_rand | 3381 | 1606 | 2905 | 1218 | 2662 |
| Activated_NT | 1676 | 818 | 1251 | 251 | 665 |
| Resting_NT | 1395 | 747 | 1203 | 266 | 727 |
| Activated *NUP62* KO | 1174 | 614 | 933 | 167 | 485 |
| Resting *NUP62* KO | 1041 | 589 | 842 | 224 | 588 |

| | **Lamina** | **NaN** | **Near_Lm1** | **Near_Lm2** | **Unassigned** | **Speckle** |
|---|---|---|---|---|---|---|
| Activated_RMC_covariate | 7900 | 76 | 3742 | 4805 | 2705 | 6686 |
| Activated_RMC_rand | 8859 | 712 | 6268 | 7466 | 4029 | 3043 |
| Resting_RMC_covariate | 8258 | 76 | 3987 | 4863 | 2581 | 6295 |
| Resting_RMC_rand | 8907 | 717 | 6267 | 7444 | 4034 | 3057 |
| Activated_NT | 1160 | 60 | 1193 | 1538 | 455 | 3361 |
| Resting_NT | 1643 | 108 | 1240 | 1733 | 410 | 2381 |
| Activated *NUP62* KO | 828 | 81 | 850 | 1038 | 272 | 2214 |
| Resting *NUP62* KO | 1525 | 127 | 1020 | 1586 | 307 | 1721 |
